# Desert lizard diversity worldwide: effects of environment, time, and evolutionary rate

**DOI:** 10.1101/2021.02.09.430433

**Authors:** Héctor Tejero-Cicuéndez, Pedro Tarroso, Salvador Carranza, Daniel Rabosky

**Affiliations:** Institute of Evolutionary Biology (CSIC-Universitat Pompeu Fabra), Barcelona, Spain; CIBIO/InBIO, Research Centre in Biodiversity and Genetic Resources, Universidade do Porto, Portugal; Museum of Zoology, Department of Ecology & Evolutionary Biology, University of Michigan, Ann Arbor, MI 48109

**Keywords:** Arid regions, regional diversity, species richness, diversity anomalies, evolutionary time, speciation rate

## Abstract

**Aim:** Biodiversity is not uniformly distributed across the Earth’s surface, even among physiographically comparable biomes in different biogeographic regions. For lizards, the world’s large desert regions are characterized by extreme heterogeneity in species richness, spanning some of the most species-rich (arid Australia) and species-poor (central Asia) biomes overall. Regional differences in species diversity may arise as a consequence of the interplay of several factors (e.g., evolutionary time, diversification rate, environment), but their relative importance for biogeographic patterns remains poorly known. Here we use distributional and phylogenetic data to assess the evolutionary and ecological drivers of large-scale variation in desert lizard diversity.

**Location:** Deserts worldwide.

**Major taxa studied:** Lizards (non-snake squamates).

**Methods:** We specifically test whether diversity patterns are best explained by differences in the ages of arid-adapted lineages (evolutionary time hypothesis), by regional variation in speciation rate, by geographic area of the arid systems, and by spatial variation related to environment (climate, topography, and productivity).

**Results:** We found no effect of recent speciation rate and geographic area on differences in desert lizard diversity. We demonstrate that the extreme species richness of the Australian deserts cannot be explained by greater evolutionary time, because species began accumulating more recently there than in more species-poor arid regions. We found limited support for relationships between regional lizard richness and environmental variables, but these effects were inconsistent across deserts, showing a differential role of the environment in shaping the lizard diversity in different arid regions.

**Main conclusions:** Our results provide evidence against several classic hypotheses for interregional variation in species richness, but also highlight the complexity of processes underlying vertebrate community richness in the world’s great arid systems.

## INTRODUCTION

What causes the dramatic differences in species diversity that we observe among different regions of the world? The latitudinal diversity gradient (LDG) is perhaps the most famous spatial diversity pattern, but species richness can vary greatly even among physiographically similar but geographically distinct biomes (Couvreur, 2015; Kelt et al., 1996; Pianka, 1986; Ricklefs & Schluter, 1993; Valente & Vargas, 2013). There is overall agreement that multiple ecological and evolutionary factors influence the assembly of regional species pools through their effects on dispersal, speciation, and extinction (Cracraft, 1992; Rabosky, 2009; Ricklefs, 1987; Weber, Wagner, Best, Harmon, & Matthews, 2017; Wiens, 2011), but we still lack general consensus on the relative importance of processes that shape regional differences in species richness (Belmaker & Jetz, 2015; Mittelbach et al., 2007).

A key challenge with large-scale assessments of species richness is that climatic, historical, and geographic hypotheses for species diversity may covary, such that disentangling their relative contributions to species diversity may be difficult or impossible (Etienne et al., 2019; Rabosky & Hurlbert, 2015). For example, in the case of the LDG, the tropics are generally older, larger, warmer, and more productive than extratropical regions. These factors are generally predicted to affect species richness in the same direction, and it is therefore difficult to isolate the effects of any single process on the overall LDG. In contrast, regional differences in species richness among climatically-matched regions can facilitate tests of the role of historical and geographic factors on diversity, by controlling – at least in part – for broad-scale differences in climate between regions. Many previous studies have addressed the role of evolutionary processes in generating these “diversity anomalies” (Ricklefs & Latham, 1993) through intercontinental comparisons of tropical rainforests (Couvreur, 2015), temperate floras (Latham & Ricklefs, 1993; Qian & Ricklefs, 2000), Mediterranean floras (Valente & Vargas, 2013), mangrove ecosystems (Ricklefs & Latham, 1993), and others.

One of the most striking regional diversity anomalies is the exceptional species richness of Australian desert lizards relative to lizards from other deserts (Pianka, 1969, 1971, 1972). Arid Australia has long been known to have outstanding lizard diversity at the biogeographic scale (i.e., the total number of species in the arid zone) compared to other arid systems of the world (Pianka, 1985, 1986). In a recent synthesis of global reptile distributions, (Roll et al., 2017) found strong support for exceptional regional diversity (richness in one-degree grid cells) of lizards in arid Australia. This diversity holds not only in comparison to other arid systems, but even in comparison to lowland tropical rainforests (Roll et al., 2017). At the local scale, more than 70 species of lizards (non-snakes) can occur in regional sympatry in arid Australia, a number that far exceeds those for the world’s second-most species-rich lizard communities (e.g., Amazon rainforest, with 25-35 species; Duellman, 2005; Rabosky et al., 2019). Numerous hypotheses have been proposed to explain these dramatic differences in the number of species, including vegetative structure, habitat specificity and heterogeneity, termite abundance, and other ecological, biogeographic, and historical factors (Morton & James, 1988; Pianka, 1972, 1989).

The outstanding richness of Australian lizards at local, regional, and biogeographic scales (Pianka, 1969, 1986), suggests the possibility that ecological and evolutionary factors cascade upwards and/or downwards to influence species richness at larger (or smaller) spatial scales. For instance, analysing range size and overlap for the diverse (∼100 species) Australian lizard genus *Ctenotus*, James & Shine (2000) concluded that the high local species richness might be related to the large area of the arid zone, and the high regional diversity can be explained by the wide distribution ranges favoured by climate homogeneity. Thus, local richness may, at least in part, reflect processes that contribute to overall high species diversity within the biogeographic and regional species pools (Ricklefs, 1987, 2004), such that evolutionary and historical factors operating at regional-biogeographic scales also shape species richness at local scales.

High levels of species diversity can be the outcome of different, but not mutually exclusive, factors (Jablonski, Huang, Roy, & Valentine, 2017). For example, high diversity might be simply due to the relatively ancient and undisturbed history of one region and the lineages within (evolutionary time hypothesis; Fine & Ree, 2006; Fischer, 1960; Mittelbach et al., 2007; Wallace, 1878). Net diversification rates, either through faster speciation or lower extinction, might also contribute to elevated species richness (Schluter & Pennell, 2017). Other explanations include physical characteristics of the regions, which in turn might affect species richness through their impact on evolutionary rates or through their impact on the level at which diversity is regulated by diversity-dependent factors. Large geographic areas may offer more opportunities for speciation than smaller regions (Rosenzweig & Sandlin, 1997), and climate and topography, as well as environmental heterogeneity, can further influence species diversity through impacts on primary productivity, habitat and resource availability, and diversification rate (Antão et al., 2020; Badgley et al., 2017; Cantalapiedra, Domingo, & Domingo, 2018; Garcia-Porta et al., 2019; Menéndez et al., 2021; Stein et al., 2015; Stein, Gerstner, & Kreft, 2014; Velasco et al., 2018).

Here, we address the causes of the dramatic differences in lizard species richness across desert regions, with an emphasis on the exceptional diversity in arid Australia, focusing in particular on large (regional to biogeographic) spatial scales. We deliberately omit snakes from our assessment, because geographic patterns of snake diversity are markedly divergent from those for other (non-snake) squamate reptiles (e.g., “lizards”); snake diversity is generally highest in humid tropical lowlands and is broadly congruent with patterns in vascular plants, frogs, birds, and many other taxa (Roll et al., 2017). Through targeted comparisons between the largest arid systems worldwide, we test a series of hypotheses that could explain pronounced differences in lizard species richness at the biogeographic and regional scales across these (broadly-comparable physiographic) regions. Specifically, we test whether species diversity in these systems is related to: i) the age of the arid-adapted lineages in each system (evolutionary time hypothesis), ii) within-region speciation rates, iii) geographic area of the system, iv) environmental variables that could also affect the carrying capacity of the system, as well as a measure of spatial heterogeneity for each of them: topography, mean annual temperature, mean diurnal range, annual precipitation, precipitation of the driest month, aridity, potential evapotranspiration, and productivity.

After decades of interest in these questions, the increasing availability of data and the development of new comparative phylogenetic and spatial methods allow us to finally make broad comparisons between these regions from an evolutionary perspective (Schemske & Mittelbach, 2017). Although our analyses cannot conclusively identify the factors that cause geographic differences in species richness, they can reject potential explanations (Etienne et al., 2019; Rabosky & Hurlbert, 2015).

## MATERIAL AND METHODS

### Delimitation of arid systems and species categorization

We analysed lizard faunas in the deserts of the world, understanding lizards to represent all non-snake squamate reptiles. We first identified Earth’s largest arid systems, focusing on those delineated by Olson et al. (2001) in the Biome “Deserts and Xeric Shrublands” (Biome 13). Within this biome, we considered the major regions to be those with a substantial component of arid and hyper-arid conditions, that is, with an Aridity Index (AI) < 0.2 (Parsons & Abrahams, 1994; Rocha, Godinho, Brito, & Nielsen, 2021). This delimitation set includes 8 main arid systems (Figure 1): Atacama, Australian desert, Central Asian desert, Gobi, Kalahari, North American desert, Persian desert, and Saharo-Arabian desert. These ‘main arid systems’ result from the aggregation of contiguous desert regions into single large arid zones (e.g., the North American arid system includes the Sonoran, Mojave, Chihuahuan and Great Basin arid subregions; the Saharo-Arabian system includes the Sahara desert and several arid regions through the Arabian Peninsula and the Horn of Africa; etc.). Our delimitation of contiguous arid systems (e.g., we merged the Sahara and the Arabian deserts, and kept the Central Asian and the Persian deserts as separate entities) was based on a regionalisation analysis through the interactive online tool for delimiting biogeographical regions from species distribution Infomap Bioregions (Edler, Guedes, Zizka, Rosvall, & Antonelli, 2017), which uses previously described clustering methods (Rosvall & Bergstrom, 2008; Vilhena & Antonelli, 2015). For this analysis, we used global reptile distribution data (Roll et al., 2017; see below), setting the cluster cost to 1.0 and implementing 10 trials (Edler et al., 2017).

**Figure 1.**
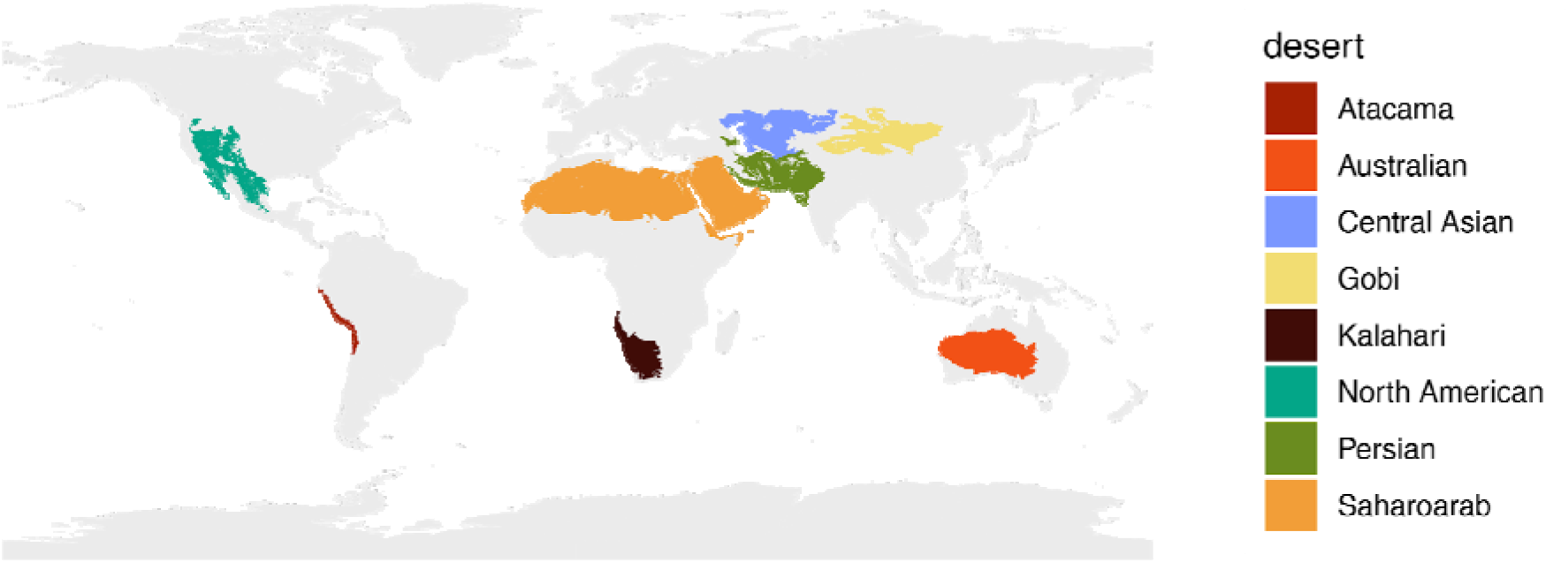
Desert systems considered in this study. We focused on regions with aridity indices (AI) < 0.2, which describe systems that are formally “arid” or “hyper-arid” but which exclude “semi-arid” (Parsons and Abrahams, 1994; Rocha et al., 2021). Map in an equal-area Behrmann projection.

We used lizard distribution data from a recently published global reptile dataset (Roll et al., 2017) to categorize each lizard species into arid systems, following twofold criteria: a species was considered as inhabiting a desert when at least 15% of its geographic range falls within the desert, or when at least 15% of the total desert’s geographic area is occupied by the species. We tested the sensitivity of the categorization percentage across a range of thresholds, and we found that the assigned species (and total diversities) for each desert were similar across a broad range of numerical thresholds (Figure S1).

### Evolutionary time hypothesis

The simplest hypothesis to explain geographic differences in species diversity relates to the total evolutionary time over which species have accumulated in different regions. This idea dates back to Wallace (1878) who proposed that the tropical diversity represents a ‘more ancient world’ than the temperate zones to explain the latitudinal diversity gradient. In order to test this hypothesis, which we refer to as the ‘evolutionary time hypothesis’, we estimated the times when lineages started diversifying within the different arid systems of the world. To do that, we reconstructed the ancestral presence of squamates in the deserts after assigning extant species to arid systems. We used the consensus tree of squamates from Tonini et al. (2016), excluding all the species without genetic or distribution data. The inclusion of species whose phylogenetic position is imputed from taxonomic data alone leads to predictable biases in ancestral state and evolutionary rate estimation and should be avoided (Rabosky, 2015). The resulting tree contained 5,355 (out of a total of 11,183; Uetz, Freed, & Hošek, 2021) squamate species (3,828 lizards and 1,526 snakes, plus the outgroup of all squamates, the tuatara; Table S1 shows the percentage of squamates, lizards, and snakes of this pruned tree in each desert relative to the phylogeny of all squamates). We assigned each one to the arid systems or categorised as “non-arid”. This classification resulted in 38 species that were assigned to more than one desert, distributed in two or more contiguous systems: Gobi, Central Asian, Persian, and Saharo-Arabian deserts (Table S2). We applied a model of character evolution with each species coded as one out of 14 possible states: “non-arid”, “North American”, “Central Asian”, “Central Asian + Gobi”, “Central Asian + Gobi + Persian”, “Central Asian + Persian”, “Central Asian + Persian + Saharo-Arabian”, “Atacama”, “Australian”, “Gobi”, “Kalahari”, “Persian”, “Persian + Saharo-Arabian”, “Saharo-Arabian”. For the ancestral state estimation we used the function ‘make.simmap’ from the R package ‘phytools’ (Revell, 2012), with an all-rates-different (ARD) model and 100 simulations. For phylogenetic data manipulation we used the R packages ‘geiger’ (Harmon, Weir, Brock, Glor, & Challenger, 2008) and ‘treeio’ (Wang et al., 2020). This approach allowed us to estimate the age at which lizard lineages started accumulating in each arid system. Critically, we do not interpret character state changes to imply biogeographic movements out of or into a particular desert. For example, an apparent transition from “Australian” to “non-arid” could simply reflect an evolved shift in biome for a lineage. Such a shift could occur even if species ranges are static through time: for example, a biome might expand or contract through time, and species might simply adapt to the new climate regime with little or no geographic movement (Donoghue & Edwards, 2014). This is particularly relevant for our analysis of ‘Desert colonisation dynamics’ (see below).

We estimated the general trajectory of lineage accumulation for each arid zone by summing the cumulative ancestral state probabilities for a given region through time (Mahler, Revell, Glor, & Losos, 2010). This metric is clearly biased by the fact that only lineages with extant descendants contribute to the lineage totals at each point in time, and thus cannot detect earlier radiations within arid regions that failed to leave extant descendants. However, in our case, we are testing hypotheses to explain the observed phylogenetic structure of arid-zone lizard diversity, and thus it is the lineage accumulation curve for extant species that is most relevant. If evolutionary time is the dominant factor influencing species richness, we expect the lineage accumulation curve for the most species-rich deserts (e.g., Australia) to predate the curves for other major arid regions. We performed correlation tests between the total number of species in each desert and the estimated time by which each desert had accumulated 10% or 30% of their total lineage diversity in the ancestral reconstruction, as a way of testing if faunal build-up started earlier in high-diversity deserts. We note that a positive correlation between age and richness is not diagnostic for identifying a contribution of clade- or regional age on species diversity. Even if diversity in each region is strictly equilibrial (and thus, not controlled by age), more diverse regions can appear “older”, simply because time to common ancestry for a regional community in equilibrium models should be positively correlated with the equilibrium number of species (Rabosky & Hurlbert, 2015). However, a negative correlation does reject the effect of age on richness. If a species-rich biota is found to be “young”, it is certainly possible that an unobserved history of diversification predates the observed lineage accumulation curve (Louca & Pennell, 2020; Quental & Marshall, 2010; Rabosky, 2016). However, it cannot then be the case that age itself is the control on richness, because the observed “young” age would have still been fully sufficient to account for all extant species richness in the species-rich region.

### Desert colonisation dynamics

Colonisation (immigration) is a key process that affects regional biodiversity patterns, along with evolutionary processes of speciation and extinction (Crouch, Capurucho, Hackett, & Bates, 2019; Hua & Bromham, 2020; MacArthur & Wilson, 1967; Simberloff, 1974). We explored the dynamics of colonisation of each desert, as well as the lizard interchange between contiguous desert regions. With such purpose, we extracted the mean number of biome shifts from external (non-arid) biomes to each arid system, as well as between arid systems. This information resulted from the 100 simulations of the ancestral reconstruction of desert occupancy that we implemented to test the evolutionary time hypothesis (see above).

### Species richness, speciation rates and geographic area of the deserts

Another evolutionary mechanism that might explain richness patterns is variation in diversification rates, e.g., areas and clades with higher diversification rates will presumably have higher species diversity. In light of difficulties in estimating extinction rates from molecular phylogenies (Rabosky, 2016), we tested this hypothesis using recent speciation rates only (tip rates), which is likely the parameter that can be most reliably inferred from a time-calibrated phylogeny (Louca & Pennell, 2020). All else being equal, elevated speciation for a given biome should translate into elevated net diversification, unless extinction rates are also sufficiently increased as to offset the effects of increased speciation rates. We estimated recent speciation rates using the DR metric (Jetz, Thomas, Joy, Hartmann, & Mooers, 2012). Although sometimes presented as an estimate of net diversification rate, DR is a much better estimate of speciation rate (Title & Rabosky, 2019). Using the Tonini et al. (2016) pseudoposterior distribution of fully-sampled squamate phylogenies, we calculated the mean DR metric for each of the 9,755 species of squamates (lizards and snakes) across 100 trees from the posterior. The Tonini et al. (2016) phylogenies are complete at the species level but include approximately 4,000 species whose position was imputed from taxonomic information using a stochastic birth-death polytomy resolver (Thomas et al., 2013). DR values computed from such trees are relatively robust to biases caused by incomplete taxon sampling and afford greater power to infer variation in evolutionary rates relative to estimates that use formal sampling fractions (Chang, Rabosky, & Alfaro, 2020). We then conducted nonparametric Kruskal-Wallis and Wilcoxon post-hoc analyses to compare lizard speciation rates among deserts to address how this metric relates with biogeographic species richness (the total number of species within each desert). We performed a correlation test between the log-transformed values of total number of species and mean DR values for each desert.

We also explored the relationship between species richness, speciation rates, and the geographic area of the arid systems, since the size of an area may affect both the species diversity and the speciation rates (Connor & McCoy, 1979; Rosenzweig, 1992). To do that, we carried out correlation tests between the log-transformed values of species richness and geographic area, and between mean tip speciation rates and geographic area among deserts. This allowed us to look for potential patterns to explain richness levels (e.g., highly diverse deserts with larger areas and/or faster speciation than less diverse ones). We implemented all the analyses, as well as data visualization, in the R environment version 3.6.3 (R Core Team, 2019), using the packages within ‘tidyverse’ (Wickham et al., 2019).

### Regional richness and environmental variables

We developed a desert lizard species richness grid overlaying the distribution data (Roll et al., 2017) onto an equal-area 10 arcminutes (∼18 x 18 Km; ∼0.17 degrees) resolution grid in a Behrmann equal-area projection. We believe that this resolution applied to the Roll et al. (2017) data captures the regional species pool, due to the relatively coarse nature of the geographic ranges themselves. The Roll et al. (2017) distributions are bounding curves that describe the overall regional extent of taxa but which provide no information about the local occurrence of taxa in the landscape. Hence, the community for any given point in space will include all species whose geographic ranges overlap broadly with the focal point, even if the point itself is localized to (for example) a habitat patch that is wholly unsuitable for many of the species. Furthermore, it is clear from previous work that species turnover in some of these systems would occur at a finer scale than the 18 x 18 km cells of our grid even if the Roll et al. (2017) ranges tracked local occurrences; Australia for example is characterized by a mosaic landscape with high habitat heterogeneity, and desert lizard communities show appreciable beta diversity at the scale of just a few hundred meters to several kilometres (Pianka, 1969; Rabosky, Cowan, Talaba, & Lovette, 2011). We investigated the potential influence of three types of environmental descriptors on desert lizard diversity patterns: topography, climate, and productivity. The topography data was obtained from the Shuttle Radar Topography Mission (SRTM) elevation data with 30 arcsec spatial resolution as processed by WorldClim v2.1 (Fick & Hijmans, 2017). We downloaded climate summary data for the period between 1970-2000 at 30 arcsec resolution from WorldClim v2.1 (Fick & Hijmans, 2017). We focused on climate variables that might better differentiate desert environments due to limiting effects on biodiversity. These include Annual Precipitation, Annual Temperature, Mean Diurnal Range and Precipitation of the Driest Month. Additionally, we downloaded the WorldClim data derived Aridity and Potential Evapotranspiration data from the CGIAR Consortium for Spatial Information (v.2; Trabucco & Zomer, 2019; Zomer et al., 2008). The productivity data is based on the normalized difference vegetation index (NDVI) over a period between 1981 and 2015 summarized on the NDVI3g time series (v.1; (Pinzon & Tucker, 2014). We downloaded and processed the NDVI data with the ‘gimms’ R package (Detsch, 2020) by obtaining the monthly data with 5 arcmin spatial resolution and summarizing the yearly data with maximum NDVI found. The final dataset is the average of the year maximum NDVI for the period between 1981 and 2015.

We spatially aggregated the environmental variables to match the target spatial resolution of the richness grid (10 arcmin ∼ 18 x 18 km). We used the mean and standard deviation as aggregating function and derived three variables from each downloaded environmental descriptor: 1) mean aggregated value describing the average value at each pixel in the target resolution, 2) within-pixel standard deviation describing the variation within the target pixel, which represents regional heterogeneity, and 3) neighbourhood standard deviation describing the surrounding variation to the target pixel. We chose two neighbours, equivalent to a 5×5 pixels search window at the target resolution, to produce the last variable. This variable describes how the surrounding environment might contribute to the species richness, as an additional estimation of environmental spatial heterogeneity. All spatial processing was performed with the ‘raster’ R library (Hijmans, 2020).

### Correlation between species richness and environmental descriptors

In order to quantify the relation between species richness and environmental descriptors, we used a non-parametric Kendall correlation score under a randomized sub-sampling framework. We conducted per-desert correlation tests between species richness and each of the environmental variables. For each environmental predictor, we conducted three different tests, corresponding to the three aggregation types (mean value, standard deviation within pixel, and neighbourhood standard deviation; see *Regional richness and environmental variables*). For each test, we used 250 randomly selected pixels from each desert to calculate the correlation and repeated the process 1,000 times. The advantages of this framework are two-fold: first, it allows us to compare same sample size, independently of the size of the desert; second, by randomly selecting scattered pixels over the desert area we decrease the effect of spatial autocorrelation of both species richness and the environmental descriptors on the correlation score. We note that this approach does not eliminate spatial autocorrelation, but it minimises the effect of non-independence of the neighbours resulting from it. From the set with the correlation scores from each of the 1,000 replicates, we calculated the average correlation and the respective 95% confidence interval. All analyses and data visualization were implemented with the R packages ‘raster’ (Hijmans, 2020), ‘rgdal’ (Bivand, Keitt, & Rowlingson, 2019), and the data science collection ‘tidyverse’ (Wickham et al., 2019).

## RESULTS

### Biogeographic-scale lizard diversity

Out of 6,303 global lizard species in the Roll et al. (2017) dataset, we identified 58 species from the Atacama Desert, 316 from the Australian deserts, 41 from the Central Asian desert, 26 from the Gobi, 207 from the Kalahari, 191 from the North American deserts, 133 from the Persian desert and 249 from the Saharo-Arabian desert (Figure 2a). As expected, arid Australia’s lizard richness is the highest among the deserts of the world. Diversity in the Saharo-Arabian desert and in the Kalahari is also noteworthy, with more than 200 species each. The Gobi and the Central Asian deserts have the lowest diversity overall, while the North American and Persian deserts have intermediate richness (Figure 2a).

**Figure 2.**
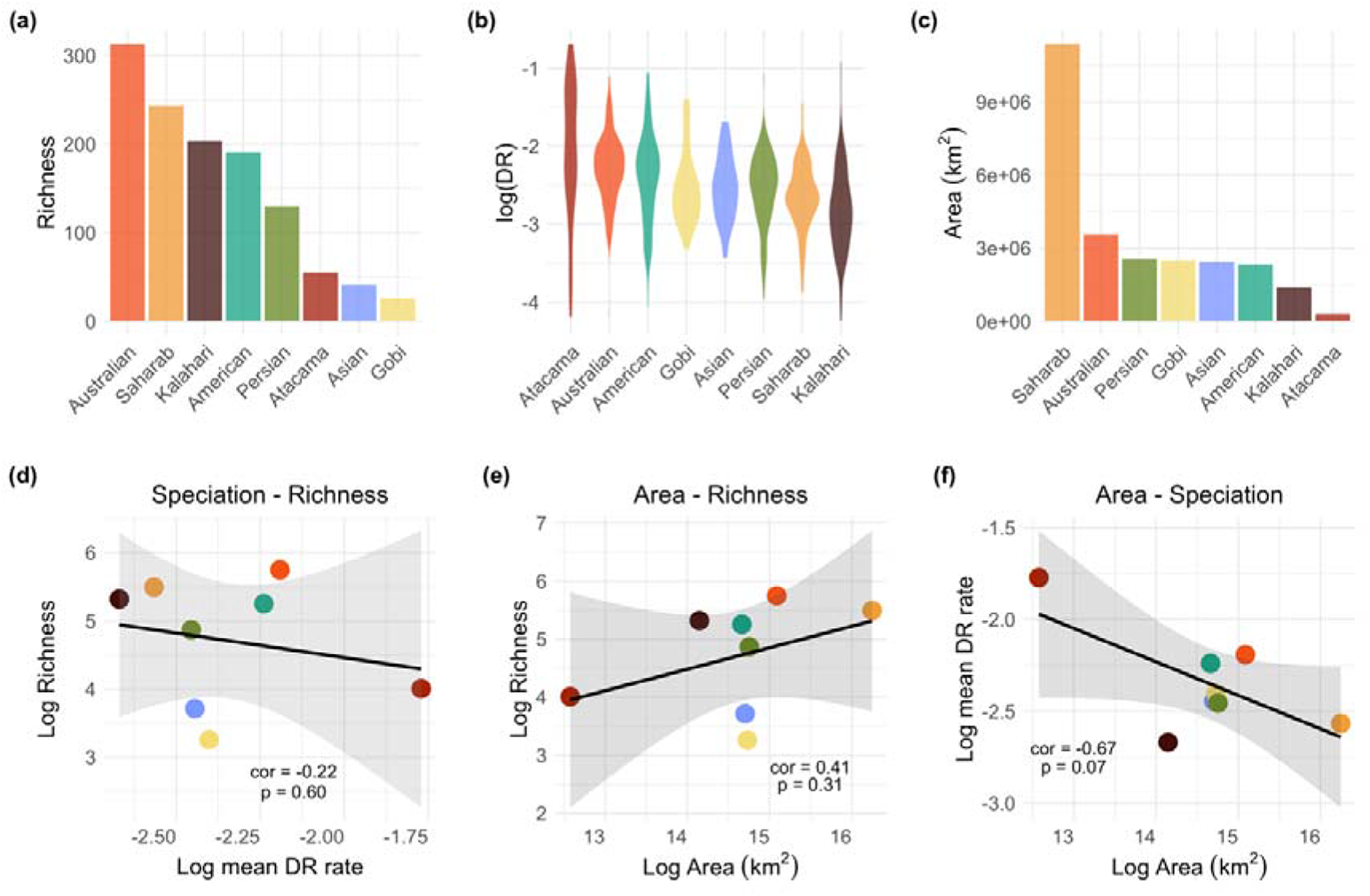
Species diversity, geographic area, and tip speciation rate across desert systems. The top row shows the characterization of arid systems: (a) biogeographic lizard richness, (b) tip speciation rates, and (c) geographic area. The bottom row shows the correlation between (d) speciation rates (mean DR) and lizard diversity in the deserts, (e) geographic area and lizard diversity, and (f) geographic area and speciation rates. There is no clear trend in the relationship between the three attributes: highly diverse deserts (e.g., Australian) are not characterized by especially high speciation rates, while several smaller and less diverse deserts (e.g., Atacama) show higher speciation rates than more diverse ones (e.g., Kalahari).

### Speciation rate and geographic area

We found significant differences in tip speciation rates (DR metric) across desert regions (Kruskal-Wallis chi-squared = 186.12, df = 7, p-value < 0.001), with highest rates in the Atacama desert and lowest in the Kalahari (Figure 2b). The post-hoc Wilcoxon test shows the pairwise differences between deserts (Table S3). We found no significant relationship between species richness and mean tip speciation rate (cor = -0.22, p = 0.60), since levels of species richness are not reflected in rate values (Figure 2a, b, d). Lizards inhabiting the most diverse deserts (Australian, Saharo-Arabian, Kalahari, American) do not show particularly high speciation rates. This decoupling is also true for systems with low lizard diversity: Atacama desert, one of the least diverse, is inhabited by taxa with the highest tip speciation rates.

We found no consistent or significant effect of geographic area on biogeographic species richness (cor = 0.41, p = 0.31) or tip speciation rates (cor = -0.67, p = 0.07; Figure 2). Regarding the richness-area relationship, the Kalahari desert, which is one of the most diverse, is the smallest one after the Atacama. In the other extreme, the Saharo-Arabian desert, of unparalleled extension, has lower diversity than Australia, and only few more species than much smaller deserts such as the North American and the Kalahari (Figure 2a, c, e). In respect of the area-speciation rate relationship, large deserts such as the Saharo-Arabian and the Australian do not present higher tip speciation rates than smaller ones such as the Atacama or the North American desert (Figure 2b, c, f).

### Evolutionary time hypothesis

The ancestral state reconstruction for desert occupancy (Figure S2) allowed us to estimate the age at which desert lizard diversity started accumulating and how the number of arid-adapted lineages has increased through time according to the reconstructed node probabilities (Figure 3a shows the rise of desert occupancy cumulative probability over time in the four most diverse deserts). Among the deserts with higher species diversity, the Saharo-Arabian and the Kalahari are the ones in which lineages started accumulating earlier, around 80-70 million years ago (Ma). Lizard diversity in the North American and the Australian deserts started increasing around 60 Ma, with North American diversity remaining at low levels until around 40-30 Ma. In contrast, arid Australian diversity shows a steep rise beginning at roughly 40 Ma, with peak rates of extant lineage accumulation at approximately 20 Ma. Thus, the lizard diversity in Australia according to our reconstructed cumulative probability through time plot does not appear before than in less diverse deserts, and nonetheless reaches unparalleled levels after such steep increase. In fact, our correlation tests yielded a negative relationship between the total richness of the deserts and the age in which lizard communities began to accumulate in each desert (e.g., the times when the ancestral state probability was a 10% or a 30% of the total cumulative probability in each desert; Figure 3b and 3c, respectively).

**Figure 3.**
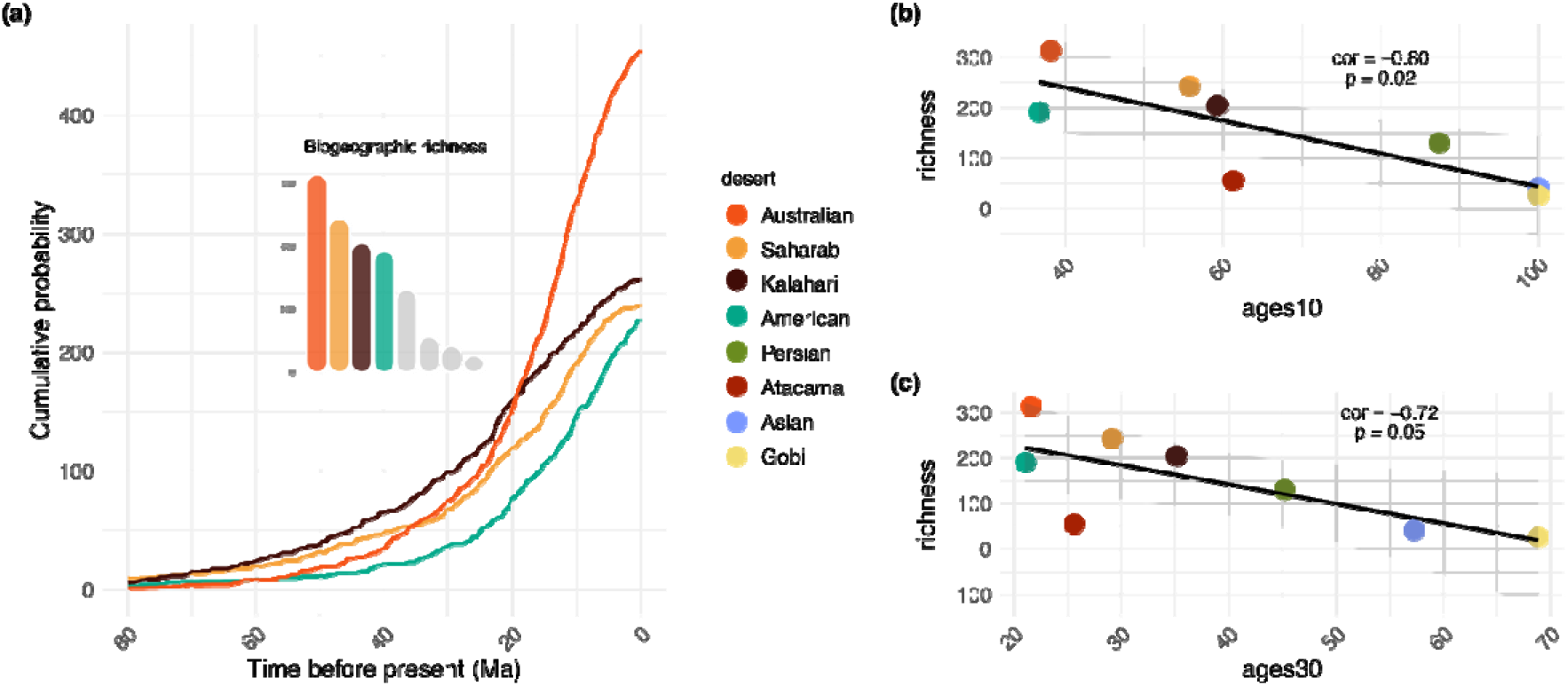
Evolutionary time and species accumulation across major desert regions. a) Reconstructed lineage accumulation curves for the four most diverse arid systems: Australian, Saharo-Arabian, Kalahari, and North American deserts. Lineage richness is represented by the cumulative probability of the focal state (arid region) summed over all nodes in the phylogeny that diverged prior to a given time point. This method necessarily underestimates true richness at a given time as it cannot accommodate extinct (unobserved) lineages, but nonetheless tracks the temporal accumulation of present-day lineages (extant) through time within regions. Panels in the right show the correlation between richness and the time in which each desert had (b) the 10% (ages10) and (c) the 30% (ages30) of their total cumulative probability in the ancestral reconstruction. Remarkably, there is a negative correlation between the age in which (extant) diversity started accumulating and the current number of species in the deserts. Namely, the most-species rich region (Australia) is associated with the most recent faunal buildup, thus rejecting the hypothesis that Australia’s exceptional diversity can be explained by greater time for diversity to accumulate.

### Desert colonisation dynamics

Ancestral state reconstruction analysis of desert occupancy implies contrasting dynamics of biome shifts in high- and low-diversity regions present essentially different dynamics of biome shifts (Figure 4; Table S4). Specifically, we observed that lineages from the most species-rich deserts (Australian, Saharo-Arabian, Kalahari, North American, and Persian) have undergone shifts to non-arid climate states much more frequently than they have adapted to arid conditions. On the contrary, less diverse deserts show a more balanced pattern of lineages shifting from non-arid to arid biomes and vice versa (e.g., Central Asian and Gobi deserts), or even an inverse pattern where the evolution of the arid state has been the prevailing direction of change (e.g., Atacama desert). Additionally, we found a generally balanced interchange between contiguous desert regions (e.g., between the Saharo-Arabian and the Persian, and between the Persian and the Central Asian).

**Figure 4.**
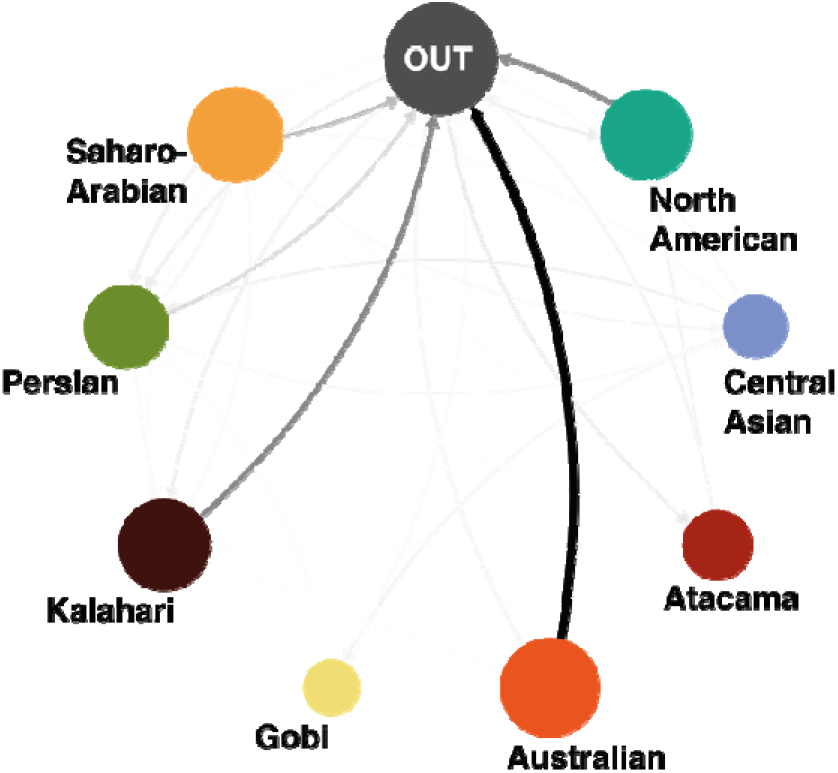
Desert colonisation dynamics: frequency of biome shifts of lineages undergoing new climate tolerances, from arid (desert systems) to non-arid (“OUT”), in each desert region. Shifts in climate tolerance, as well as lineage interchange between deserts, are depicted with directional arrows. Arrow thickness and colour intensity represent the number of events. Circle size is proportional to the number of species within each desert. Information of events into and out of the deserts are extracted from the ancestral reconstruction analysis (Figure S2; Table S4).

### Regional lizard richness and the effect of environmental variables

In our regional lizard diversity grid (10 arcmin, 18 x 18 km) we found the expected outstanding levels in Australia, with most of the arid Australian grid cells containing between 60 and 80 species (Figure 5). Regional diversity levels are medium in the North American and the Kalahari deserts, and they are low for the rest of the arid systems, especially the Gobi and the Atacama deserts, with few or no grid cells having more than 20 species. This result indicates that the moderate-to-high overall (biogeographic) species richness for the Saharo-Arabian desert (Figure 2a) is largely a result of the vastly greater area of that region and geographical turnover. After controlling for sampling area (Figure 5), it is clear that regional assemblages in Saharo-Arabia contain far fewer species than arid Australia and the Kalahari.

**Figure 5.**
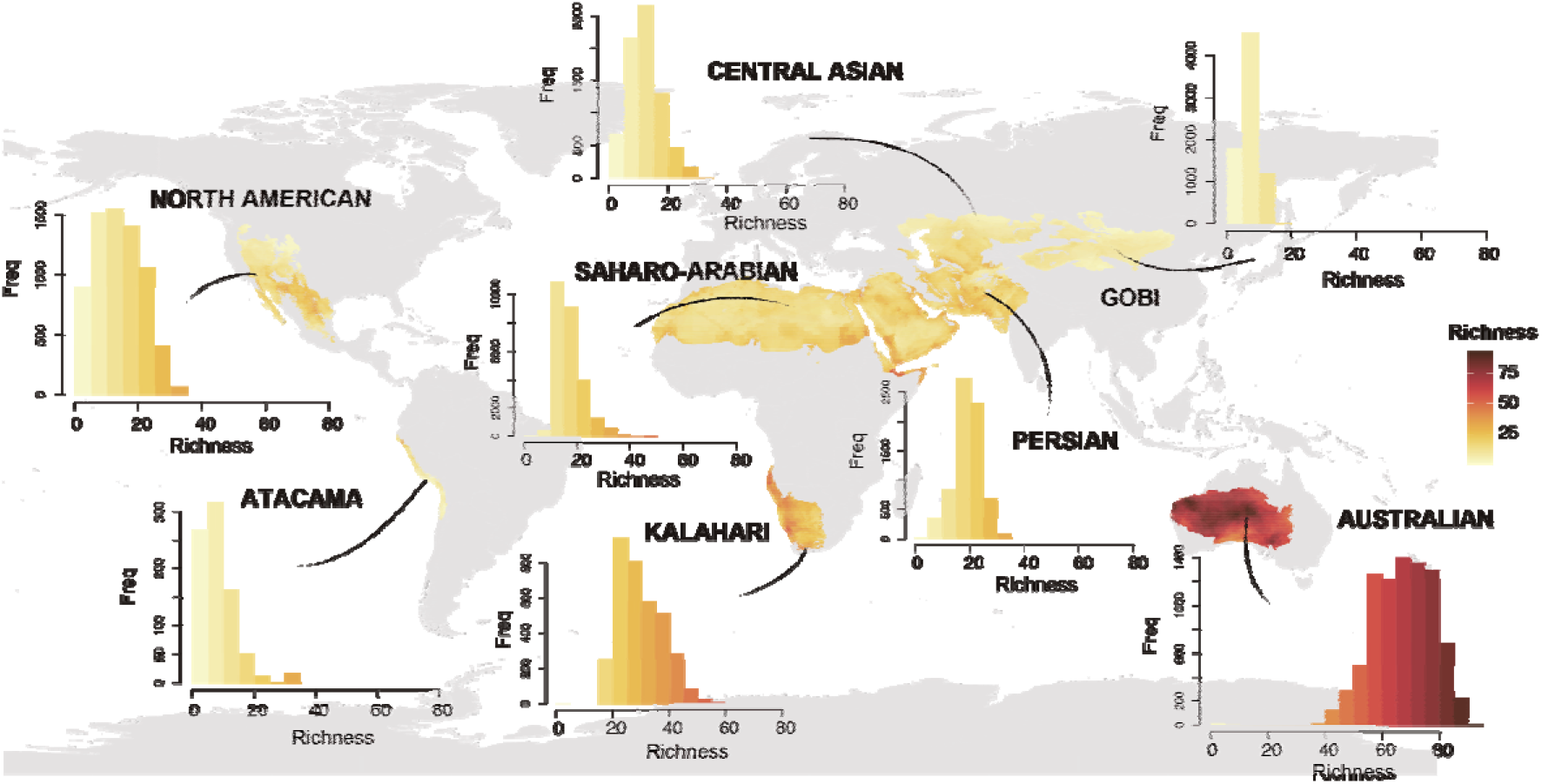
Regional lizard species richness (10 arc-minute grid cell) for the deserts of the world, with histograms of grid-cell richness for each system. Species richness of regional lizard assemblages from Australia is far greater than in any other arid region, and the system is thus exceptionally diverse at local (Pianka, 1969), regional, and biogeographic (Figure 2) scales. Map in an equal-area Behrmann projection.

We found no consistent effects of the environmental descriptors or aggregation types on regional (grid-cell) assemblages within individual deserts (Figure 6; Figure S3; Table S5, Appendix 1). The most notable relationships included a positive correlation of average potential evapotranspiration with lizard richness in the Central Asian and North American deserts (Kendall’s Tau 0.67±0.04 and 0.60±0.06, respectively); a positive correlation of average annual temperature in the Central Asian and the Gobi deserts (Kendall’s Tau 0.64±0.05 and 0.63±0.05, respectively); and a negative correlation of average precipitation of the driest month in the Central Asian desert (Kendall’s Tau −0.61±0.05) (Figure 6; Table S5; Appendix 1). Overall, the most noticeable pattern is the high disparity of the effects of each environmental variable in different deserts, which shows differential roles in each region.

**Figure 6.**
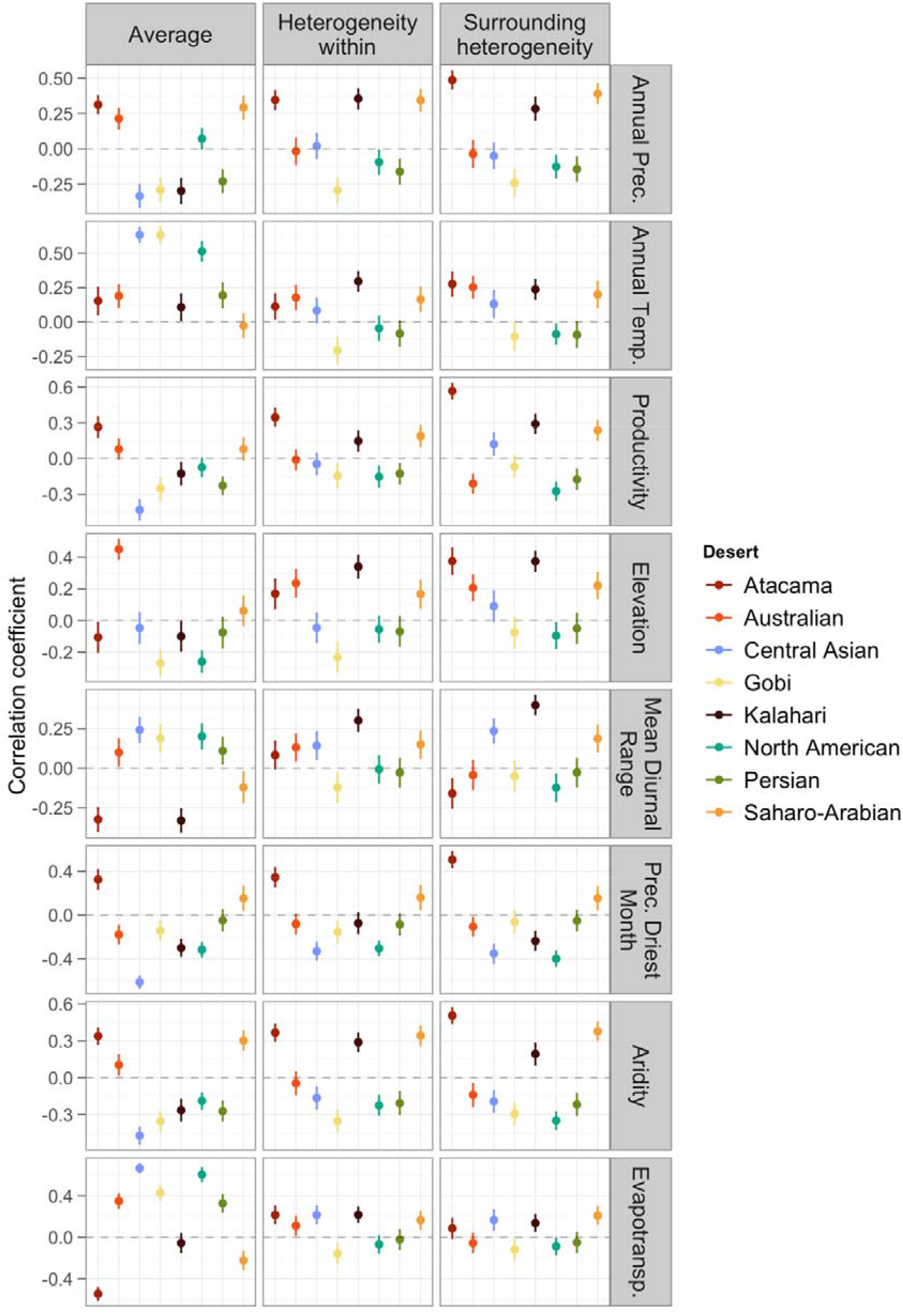
Correlation coefficient (mean and 95% confidence interval from 1,000 bootstrap replicates) between each environmental descriptor and regional lizard richness, for each arid region separately. “Average” column gives the correlations between species richness and the mean of the environmental variable per cell. “Heterogeneity within” and “Surrounding heterogeneity” give the correlation between species richness and two measures of variability in environmental conditions per cell: the standard deviation of the variable within the cell, and the standard deviation of the variable in the surrounding cells, respectively. There are no consistent effects of the environmental variables across deserts, and no clear relationships between environment and richness, although some variables seem relevant in the shaping of lizard diversity in certain deserts.

## DISCUSSION

We investigated long-standing hypotheses about the factors driving desert lizard diversity. Even though an unknown number of species remain to be described and there are inevitable differences in the knowledge of biodiversity among regions that could bias our results (Hughes et al., 2021; Meiri, 2016), we included the most recent available data on reptile distributions, finding the overall same diversity patterns that have been reported for decades.

Our results corroborate the uneven distribution of lizards across the deserts of the world and further highlight the exceptional lizard richness reported previously for arid Australia (Pianka, 1969, 1986; Roll et al., 2017), and illustrate how this pattern holds at both regional (Figure 5) and biogeographic (Figure 2) scales. Australia represents one extreme of the lizard diversity continuum among the deserts of the world, where we find some systems with medium levels of biogeographic and regional diversity, such as the North American or the Kalahari deserts, and very low levels in the other extreme, represented by deserts such as the Gobi and the Atacama. There are also cases of contrasting richness levels at different scales: the Saharo-Arabian region harbours high richness at the biogeographic scale but is relatively depauperate at the regional (grid-cell) scale. However, our results do not corroborate some current hypotheses explaining species diversity patterns in the deserts. Specifically, we find no evidence that recent speciation rates are fastest in the most species-rich deserts (Figure 2) and no evidence that the most species-rich deserts have accumulated species for longer (evolutionary time hypothesis; Figure 3). Furthermore, we found little evidence that the differences in species richness among deserts at biogeographic and regional scales are the result of the many environmental factors we considered (Figure 6). Rather than supporting universal explanations for which environmental variables determine patterns of lizard richness in the deserts, our results suggest that some of these factors might have been relevant to shape lizard diversity within the deserts, but with a differential role amongst the arid systems.

Speciation is one of the main processes that directly influence the number of species at large scales (Jetz, Thomas, Joy, Hartmann, & Mooers, 2012; Ricklefs, 1987; Schluter & Pennell, 2017). The idea of an immediate relationship between speciation rate and species richness (i.e., high levels of species richness are primarily the result of high speciation rates) is a long-standing hypothesis (MacArthur, 1969; Mittelbach et al., 2007) and has proven useful to explain the latitudinal diversity gradient in mammals (Rolland, Condamine, Jiguet, & Morlon, 2014), but it has been recently challenged in global analyses of other major groups such as fishes (Rabosky et al., 2018), birds (Jetz, Thomas, Joy, Hartmann, & Mooers, 2012; Kennedy et al., 2014; Quintero & Jetz, 2018; Rabosky, Title, & Huang, 2015), and plants (Igea & Tanentzap, 2020). The “faster speciation” hypothesis was proposed by (Pianka, 1972) to explain Australia’s extreme lizard diversity, but our study represents the first formal test of this idea. We found no relationship between high species richness and high tip speciation rates, suggesting that the exceptionally high diversity in arid Australia is not driven by faster speciation, at least by recent levels of speciation. Likewise, deserts with low species richness (e.g., Gobi and Central Asian) did not yield particularly low tip speciation rates. In fact, recent speciation can sometimes be faster in the regions with lowest diversity (Rabosky et al., 2018). Such decoupling might reflect the greater importance of extinction or immigration relative to speciation (Gaston & Blackburn, 1996; López-Estrada, Sanmartín, García-París, & Zaldívar-Riverón, 2019; Meseguer & Condamine, 2020), and/or the importance of differential carrying capacities and diversity-dependence in mediating overall diversity patterns (Marshall, 2007; Mittelbach et al., 2007; Rabosky, 2009).

It is important to note that our analyses concern present-day speciation rates only and we did not attempt to estimate net diversification rates. Our results on lineage accumulation might suggest effectively increased net diversification rates in the Australian deserts relative to other less diverse deserts. However, net diversification rates require estimates of extinction, which is notoriously difficult in the absence of fossils (Rabosky, 2016). Moreover, we did not attempt to reconstruct historical speciation rates, which might have been faster in the past; recent work has emphasized historical (deep-time) reconstructions of speciation rate from phylogenetic trees may be unreliable regardless of the inference method used to estimate them (Louca & Pennell, 2020). Nonetheless, analyses of recent speciation rates can still provide great insight into the causes of diversity gradients. Processes underlying patterns of species richness can be affected by mechanisms that are time- and diversity-independent. For example, faster speciation could result from the kinetic effects of temperature on processes that favour reproductive isolation (e.g., effects on metabolism; Allen, Brown, & Gillooly, 2002; Rohde, 1992) and as such, low-latitude deserts should be characterized by faster speciation rates than higher-latitude deserts. The shifting sandridge hypothesis of (Pianka, 1972) proposes that the mobile dune sheets within interior Australia serve as a continued source of geographic isolation for non-dune taxa — in effect, creating a continued “species pump” that will result in overall high rates of speciation for the arid Australian biota. Our finding of unexceptional recent speciation rates for the Australian deserts rejects this and other species-pump models for this system. Note that this does not mean that such shifting sandridges are unimportant for speciation, merely that speciation rates as influenced by such geomorphic/landscape processes are unlikely to serve as rate-limiting controls on species richness.

Geographic area is one of the main factors that may affect broad-scale patterns of species richness (Connor & McCoy, 1979). The concept of the species-area relationship, derived from early works (Gleason, 1922, 1925) and developed later from the island biogeography theory (MacArthur & Wilson, 1967), predicts a positive effect of geographic area on both species diversity and speciation rates (Rosenzweig, 1992; Rosenzweig & Sandlin, 1997). This has been supported for island and non-island systems (Kisel & Barraclough, 2010; Losos & Schluter, 2000; L. G. Marshall, Webb, Sepkoski, & Raup, 1982; Sepkoski, 1976; Wagner, Harmon, & Seehausen, 2014), although patterns reported for some systems have likely been confounded by assumptions that speciation rates are invariant through time (Rabosky & Glor, 2010). Moreover, some authors have argued against geographic area being the determinant factor on species diversity (Chown & Gaston, 2000; Rohde, 1992, 1998). We found no significant effect of area on species richness, although the correlation was positive (Figure 2e). It is possible that geographic area contributes to species richness, but the relationship is incredibly noisy at best: the largest desert by far is the Saharo-Arabian, which contains substantially lower diversity than Australia and roughly equal diversity to the much smaller Kalahari. Conversely, the tiny Atacama desert – by far the smallest in the dataset – has greater diversity than two much larger deserts.

Alternatively, evolutionary time may generate diversity patterns in which species richness is decoupled from geographic area and other factors (Rohde, 1986; Stephens & Wiens, 2003), although some evidence supports the combined effect of area and time (Fine & Ree, 2006; Jetz & Fine, 2012). Age differences between geographic regions or biomes has long been thought to be a potential contributor to differences in species richness (Wallace, 1878), for the simple reason that older, less-disturbed geographic regions have longer periods of time for diversity to accumulate (Fischer, 1960; Mittelbach et al., 2007). Our results argue against any simple role for time in explaining the observed heterogeneity in desert lizard richness and even suggest a negative relationship between richness and age (Figure 3). Remarkably, our results suggest that Australian lizard diversity began accumulating more than ten million years after the onset of lineage accumulation in less-diverse deserts, like the Saharo-Arabian and the Kalahari (Figure 3). Thus, greater time for diversification is unlikely to account for the extreme lizard richness observed in arid Australia, and other factors are likely responsible for the steep increase in lizard diversity observed in this region (e.g., historical diversification dynamics). Note that these results do not rely on accurately inferring the true timing of squamate diversification in each region, but only on the phylogenetic age of the extant biota. For example, now-extinct lineages of squamates may have undergone extensive diversification in arid or semi-arid Australia many millions of years prior to our reconstructions (Oliver & Hugall, 2017), but it is still true that the entire extant Australian arid zone fauna was able to accumulate in a much briefer period of time.

It is worth noting that our estimation of the age of arid-adapted lineages does not aim to serve as an absolute date for desert origination. The age of the deserts considered here is a complex and contentious issue (Swezey, 2006). The onset of arid conditions might have greatly preceded the formation of many modern-day arid landscapes (Pillans, 2018; Schuster et al., 2006), and hence could potentially explain the presence of arid-adapted lineages long before the estimated geological age of the desert (Byrne et al., 2008; Carranza, Arnold, Geniez, Roca, & Mateo, 2008). Therefore, our estimation of the age of arid-adapted lineages does not aim to serve as an absolute date for desert origination, but instead as a relative time framework to compare the biotas inhabiting different desert systems.

Regarding colonisation dynamics, we found a consistent pattern distinguishing high- and low-richness deserts (Figure 4). Species-rich deserts (e.g., Australian, Saharo-Arabian, Kalahari, North American) are characterized by high levels of phylogenetic transitions to non-arid conditions compared to shifts towards arid climate tolerance. This contrasts with species-poor deserts, such as the Gobi or the Central Asian, where evolutionary shifts from non-arid to arid conditions and vice versa are balanced. Further, in the case of the Atacama, we observed a net excess of biome transitions into the region (Figure 4; Table S4). This suggests essentially different processes of community assembly in regions with high and low diversity. Based on these results, communities in high-diversity systems build through a few colonisation events with high levels of subsequent intra-desert diversification, with many lineages of those clades secondarily adapting to non-desert conditions. Less diverse deserts, on the contrary, seem to be characterized by a relatively regular lineage interchange with regions of lower aridity, without extensive radiations within the deserts. These differences might be related to intrinsic system-specific characteristics that may offer more opportunities for speciation in some regions, resulting in high diversity, while some other regions of low diversity might serve as evolutionary ‘dead-ends’, with limited opportunities for diversification within the desert.

Despite theoretical and empirical support linking climate and topography to species richness in general (Currie, 1991; Francis & Currie, 2003; Jetz & Rahbek, 2002; Kreft & Jetz, 2007; Qian, 2010; Skeels, Esquerré, & Cardillo, 2020; Wright, 1983), we found ambiguous support for the effects of environmental variables on desert lizard diversity at both regional and biogeographic scales. Our per-desert environment-richness analyses show differential effects of the environmental descriptors on lizard richness across deserts (Figure 6). We found a considerable correlation of richness in some deserts with several variables, in particular average evapotranspiration in Central Asia and North America, average annual temperature in Central Asia and the Gobi, and (negative) average precipitation of the driest month in Central Asia. Thus, the impact of the environment on the distribution of lizard diversity in the Central Asian desert seems stronger than in other arid systems. It is interesting to note the shared patterns in the Central Asian and the Gobi desert with respect to most variables. Both deserts have similar levels of diversity (very low biogeographic and regional diversity), and this might indicate a differential role of climate and other factors between high- and low-diversity systems, as also suggested by our results on desert colonisation dynamics (Figure 4). In the Atacama desert, environmental heterogeneity (e.g., surrounding heterogeneity in precipitation, productivity and aridity) appears to play a significant role in the distribution of diversity. Average elevation is more related with the higher diversity in Australia than any other variable, and its importance for Australian lizards is remarkable in comparison with other deserts, while in the Kalahari and the Atacama deserts the elevational heterogeneity seems more relevant. Additionally, in the Kalahari, heterogeneity plays a rather constant role, although without strong correlations with richness. In the Saharo-Arabian region, on the contrary, there are not remarkable effects of the variables tested on lizard distribution. It should be noticed here that we only tested recent environmental factors. It is likely that deep-time environmental dynamics (e.g., when most of the desert diversity originated and assembled) had an important role in shaping lizard communities by affecting the stability of the arid systems through time (as well as historical diversification). In fact, the impact of biome fragmentation, expansion and contraction cycles might have been particularly relevant in the deserts (Barbolini et al., 2020; deMenocal, 1995; Hernández-Fernández & Vrba, 2005).

We did not find consistent effects of climate, productivity, and topography on species richness across deserts, but it is possible that other, unmeasured environmental variables and their interactions provide an important control on broad-scale richness. Our results, hence, are consistent with a scenario in which desert lizard diversity at regional-to-biogeographic scales depends mainly on system-specific ecological dynamics. Factors such as extinction, habitat heterogeneity, ecological preferences, and ecological interactions with other taxa, might be playing a relevant role in shaping the distribution of current lizard diversity in the deserts. Our results provide evidence against several general hypotheses proposed to explain desert lizard diversity patterns (age; evolutionary rate). However, we only studied part of the evolutionary rates hypothesis (recent speciation), and we speculate that the relationship between richness and environment may not be adequately captured with our simplified set of eight explanatory variables. Ultimately, our results emphasise the complexity of evolutionary and ecological processes that determine large-scale diversity patterns. Disentangling their underlying causes remains an essential task and one that is necessary in order to face the challenges imposed by the rapid pace of climatic and other change that is disrupting ecosystems worldwide.

## DATA AVAILABILITY STATEMENT

All the data and scripts produced in this work will be made available at an online repository (e.g. GitHub), which will be updated and made public upon acceptance of the manuscript.

## CAPTIONS FOR SUPPLEMENTARY FIGURES

**Figure S1**. Number of lizard species in each arid system under different threshold values for categorization of species into deserts: 10_10, 10_20, 15_15, etc. The number before the underscore indicates the percentage of the species distribution that needs to be within a desert to be categorized into that desert, and the number after the underscore indicates the percentage of a desert’s extension that needs to be occupied by a species for the species to be categorized into that desert. If one of those criteria is met, a species is categorized into the arid system.

**Figure S2**. Ancestral reconstruction of desert occupancy on the Squamate phylogenetic tree from Tonini et al. (2016) excluding species without genetic and distribution data. Colors at the tips indicate the arid system(s) occupied by each species, and pie charts at nodes indicate the probability of occupying each desert.

**Figure S3.** Scatterplots showing the relationship between environmental variables and grid-cell lizard richness, coloured by desert region.

## CAPTIONS FOR SUPPLEMENTARY TABLES

**Table S1**. Percentage of squamate, lizard, and snake species present in the ancestral reconstruction (i.e., with genetic and distribution data) for each desert, relative to the total number of species in the phylogeny of all squamates (Tonini et al., 2016).

**Table S2**. Categorisation of squamate species with genetic (Tonini et al., 2016) and distribution data (Roll et al., 2017) into the major arid regions of the world. This was the dataset used for the ancestral reconstruction of desert occupancy (Figure S2).

**Table S3**. Significance values (p-values) of the post-hoc tests showing differences in tip speciation rates (DR metric) between desert regions.

**Table S4**. Average number of biome transitions into and out of each desert region reconstructed in the 100 simulations of the evolution of desert occupancy (Figure S2). Rows show the ancestral state, and columns show the descendant state. For example, the first row shows the frequency of shifts from non-desert conditions (“Out”) to each of the desert regions. Likewise, the first column shows the frequency of shifts from each desert to non-arid conditions (“Out”).

**Table S5**. Results from the correlation tests between environmental variables and grid-cell lizard richness in each desert. For each correlation, 1,000 replicates were implemented, each one with a random sample of 200 grid cells. The table show the mean, the standard deviation (sd), and the 95% confidence interval (low.ci and up.ci) of the correlation coefficients from each analysis.

## CAPTIONS FOR SUPPLEMENTARY APPENDIX

**Appendix 1**. Scatterplots showing the relationship between environmental variables and grid-cell lizard richness in each desert.

