## Supplementary material for "Desert lizard diversity worldwide: effects of environment, time, and evolutionary rate": Fig. S1

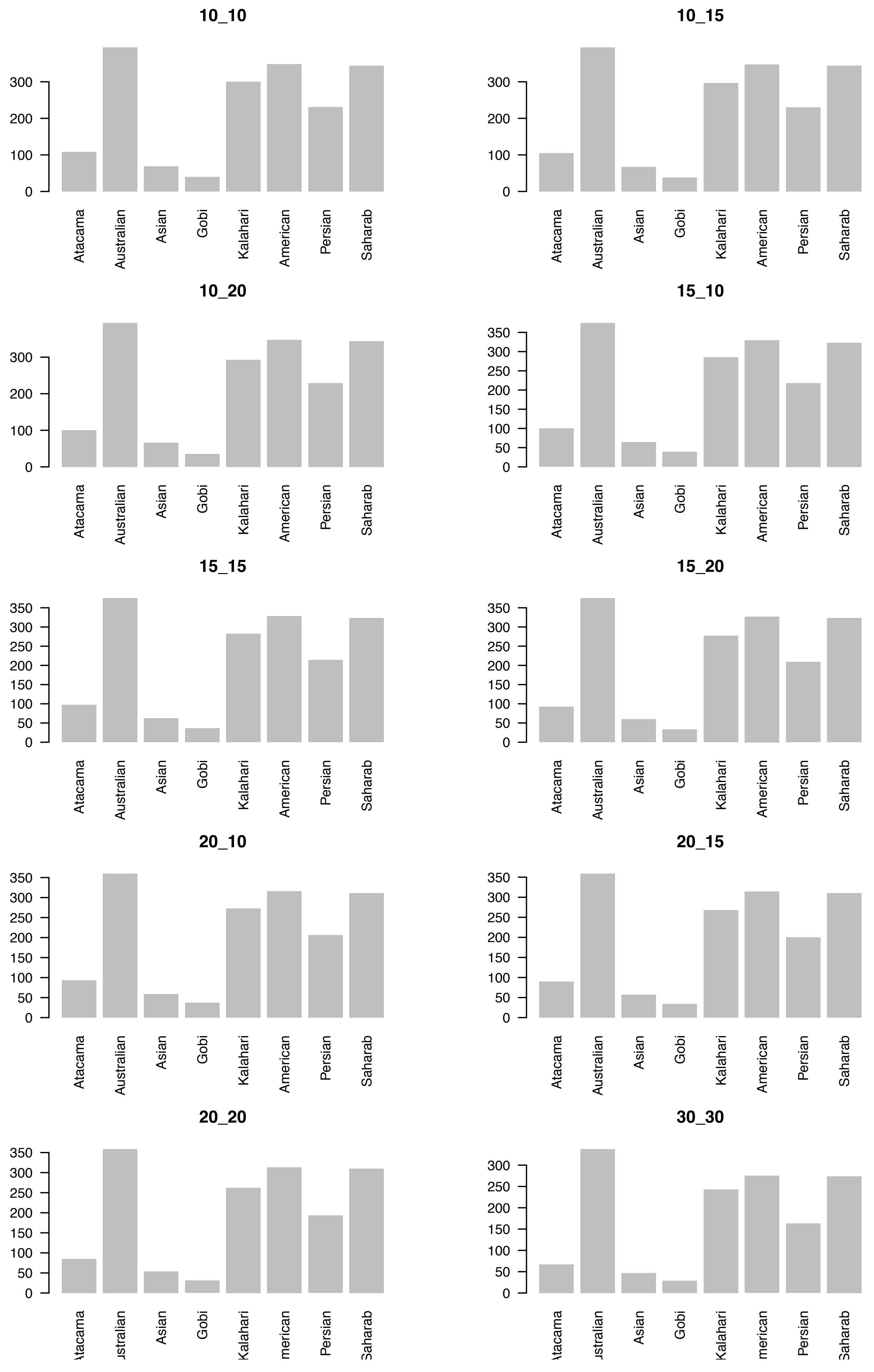


**Figure S1**. Number of lizard species in each arid system under different threshold values for categorization of species into deserts: 10_10, 10_20, 15_15, etc. The number before the underscore indicates the percentage of the species distribution that needs to be within a desert to be categorized into that desert, and the number after the underscore indicates the percentage of a desert's extension that needs to be occupied by a species for the species to be categorized into that desert. If one of those criteria is met, a species is categorized into the arid system.
