## Supplementary material for "Desert lizard diversity worldwide: effects of environment, time, and evolutionary rate": Fig. S2

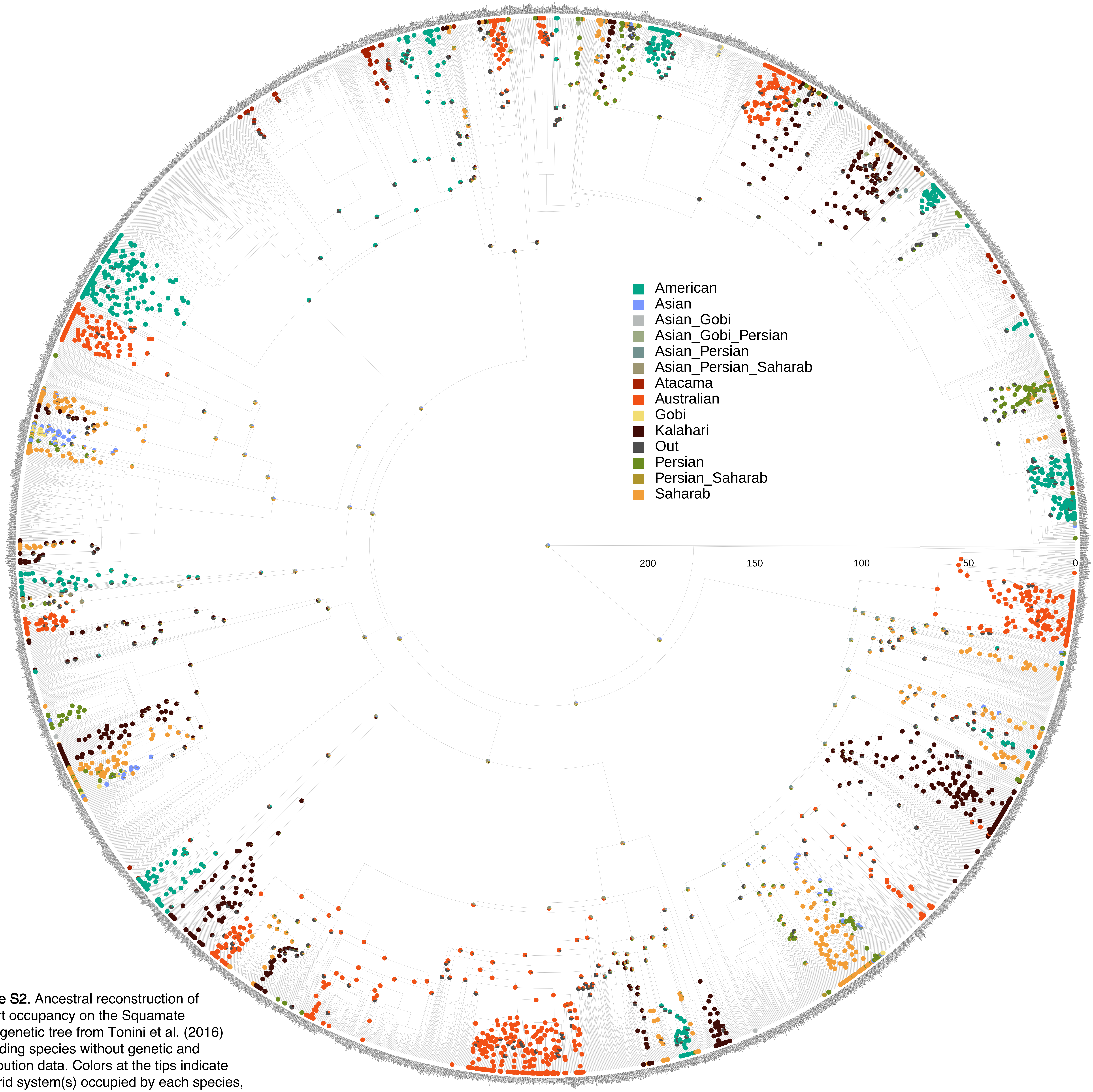


**Figure S2.** Ancestral reconstruction of desert occupancy on the Squamate phylogenetic tree from Tonini et al. (2016) excluding species without genetic and distribution data. Colors at the tips indicate the arid system(s) occupied by each species, and pie charts at nodes indicate the probability of occupying each desert.
