## Appendix 1 for "Desert lizard diversity worldwide: effects of environment, time, and evolutionary rate"

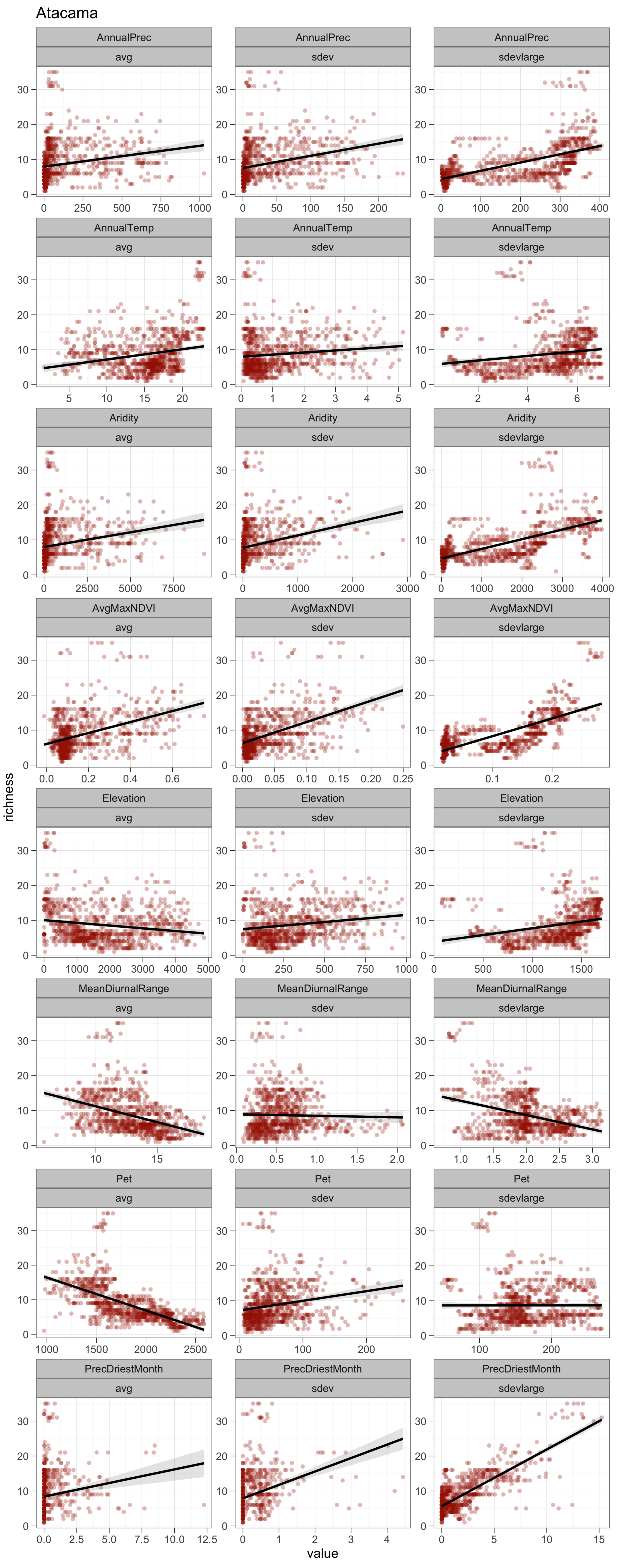
Atacama
AnnualPrec avg
AnnualPrec sdev
AnnualPrec sdevlarge
AnnualTemp avg
AnnualTemp sdev
AnnualTemp sdevlarge
Aridity avg
Aridity sdev
Aridity sdevlarge
AvgMaxNDVI avg
AvgMaxNDVI sdev
AvgMaxNDVI sdevlarge
Elevation avg
Elevation sdev
Elevation sdevlarge
MeanDiurnalRange avg
MeanDiurnalRange sdev
MeanDiurnalRange sdevlarge
Pet avg
Pet sdev
Pet sdevlarge
PrecDriestMonth avg
PrecDriestMonth sdev
PrecDriestMonth sdevlarge
richness
value

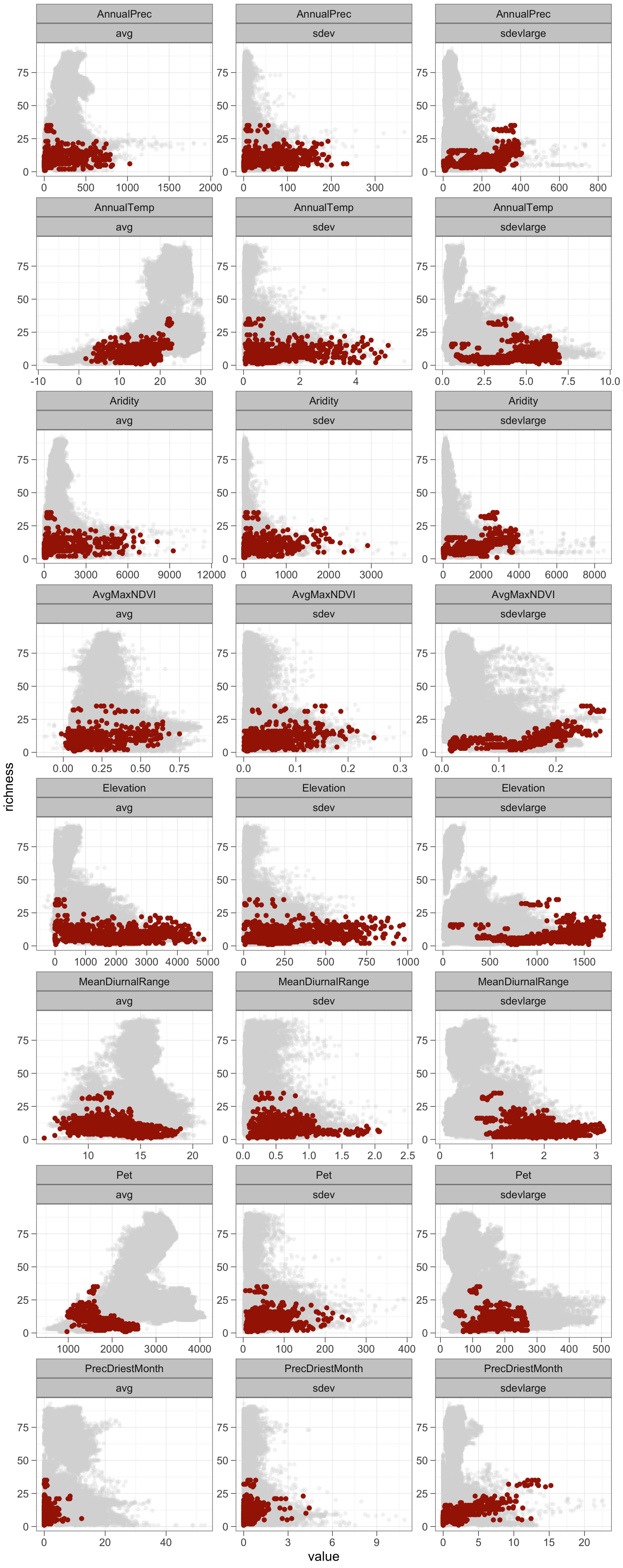

AnnualPrec
avg
sdev
sdevlarge
AnnualTemp
Aridity
AvgMaxNDVI
Elevation
MeanDiurnalRange
Pet
PrecDriestMonth
richness
value

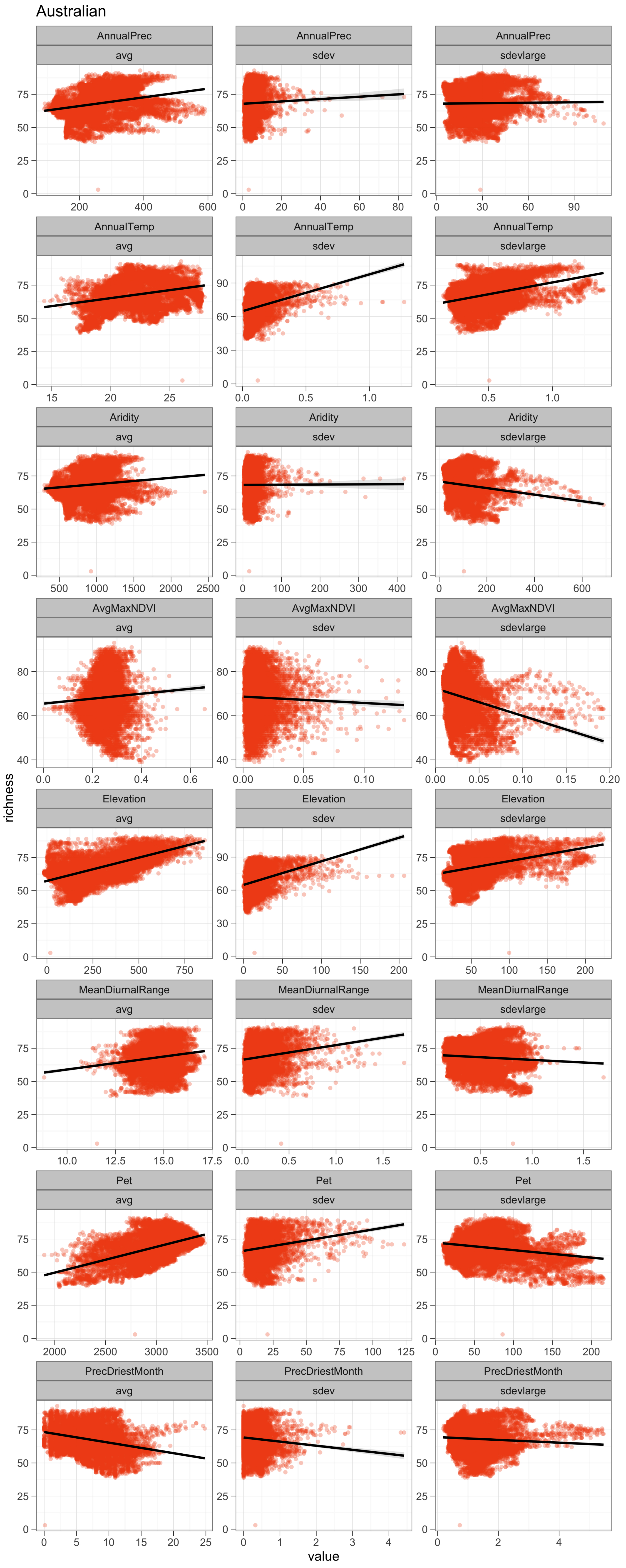
Australian
AnnualPrec
avg
AnnualPrec
sdev
AnnualPrec
sdevlarge
AnnualTemp
avg
AnnualTemp
sdev
AnnualTemp
sdevlarge
Aridity
avg
Aridity
sdev
Aridity
sdevlarge
AvgMaxNDVI
avg
AvgMaxNDVI
sdev
AvgMaxNDVI
sdevlarge
Elevation
avg
Elevation
sdev
Elevation
sdevlarge
MeanDiurnalRange
avg
MeanDiurnalRange
sdev
MeanDiurnalRange
sdevlarge
Pet
avg
Pet
sdev
Pet
sdevlarge
PrecDriestMonth
avg
PrecDriestMonth
sdev
PrecDriestMonth
sdevlarge
richness
value

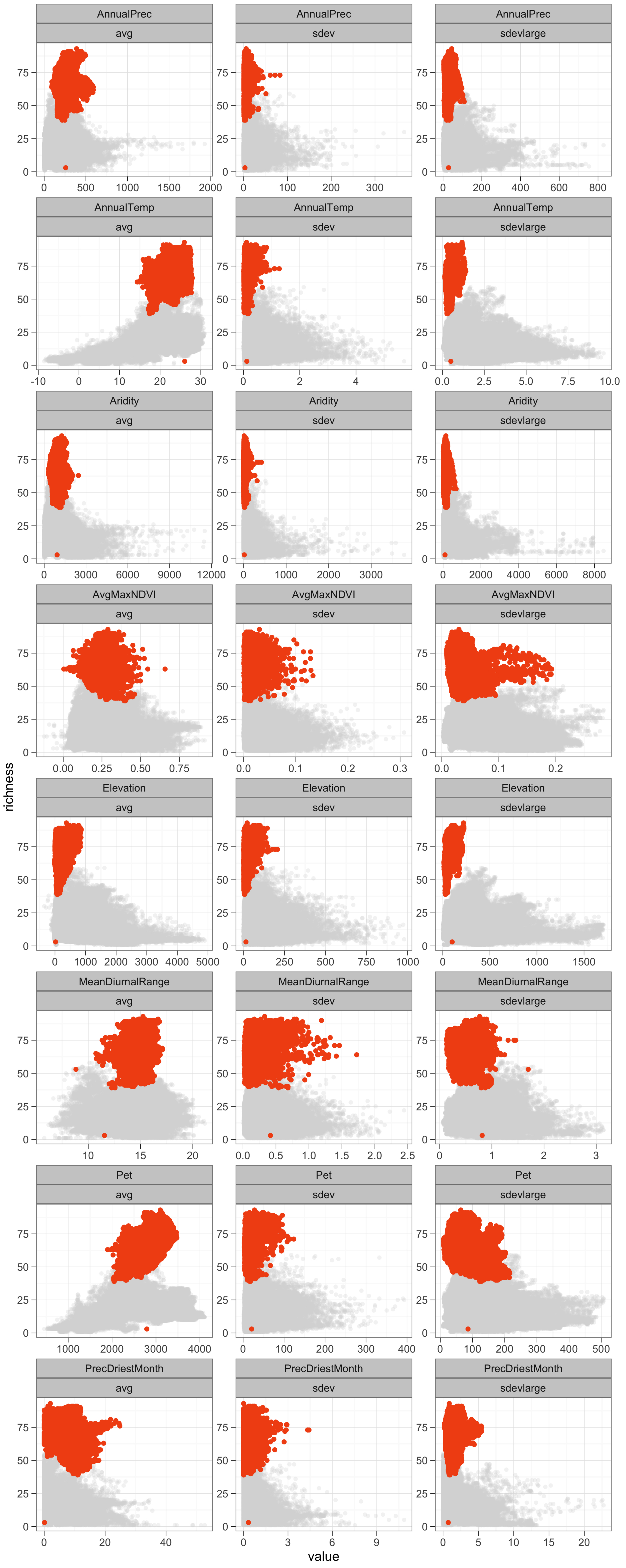

AnnualPrec
avg
sdev
sdevlarge
AnnualTemp
Aridity
AvgMaxNDVI
Elevation
MeanDiurnalRange
Pet
PrecDriestMonth
richness
value

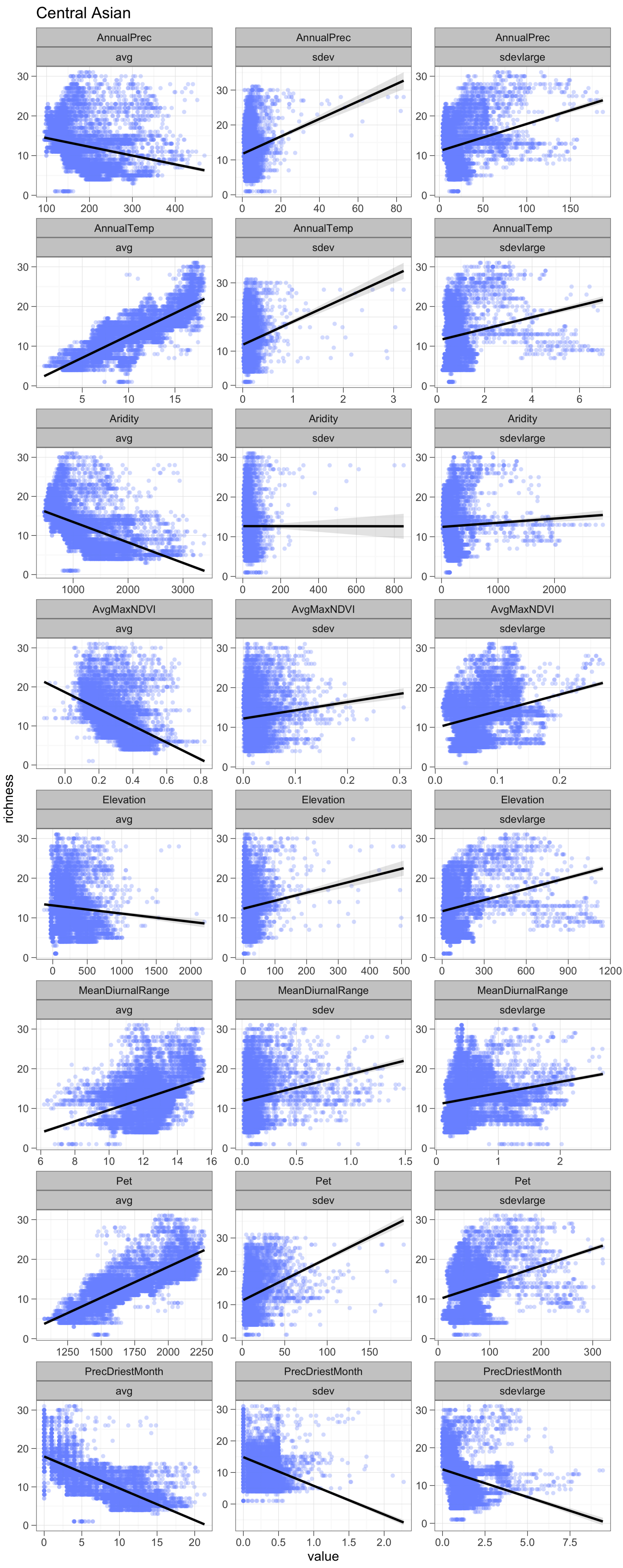

Central Asian
AnnualPrec
avg
sdev
sdevlarge
AnnualTemp
Aridity
AvgMaxNDVI
Elevation
MeanDiurnalRange
Pet
PrecDriestMonth
richness
value

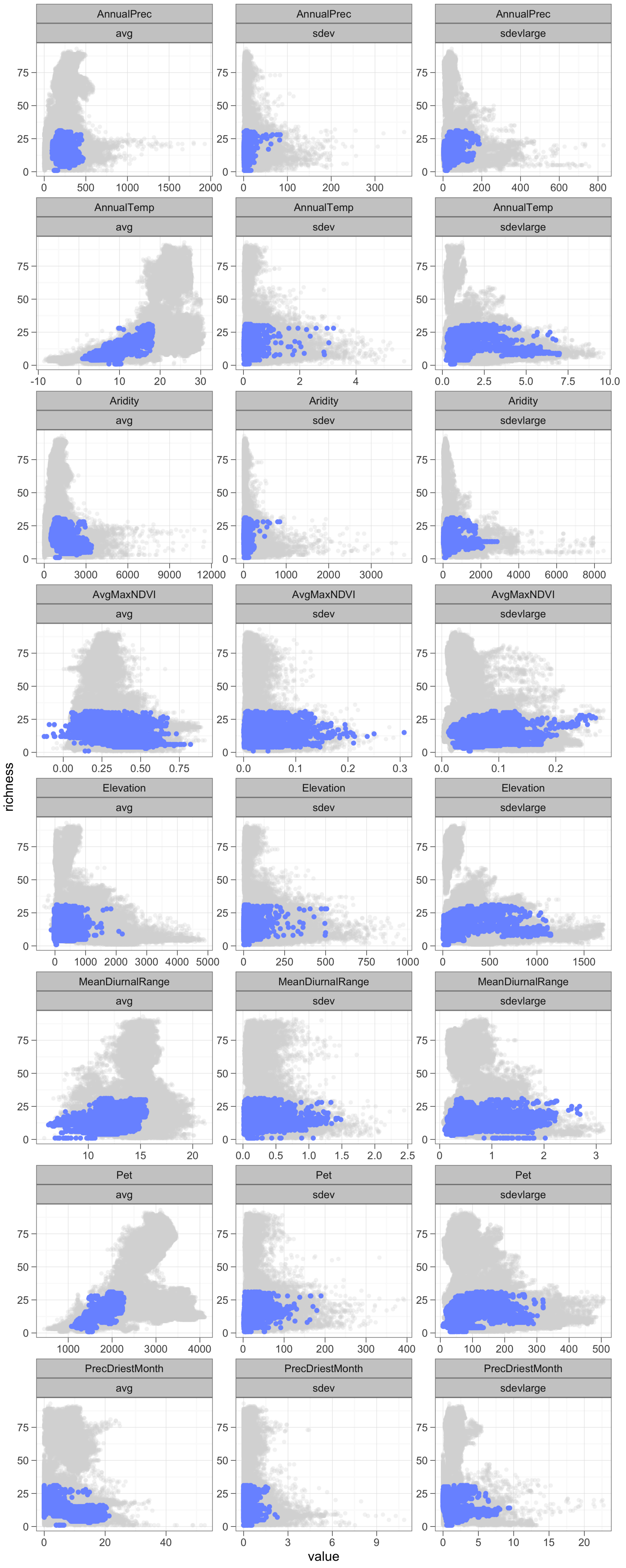

AnnualPrec
avg
0 500 1000 1500 2000
AnnualPrec
sdev
0 100 200 300
AnnualPrec
sdevlarge
0 200 400 600 800
AnnualTemp
avg
-10 0 10 20 30
AnnualTemp
sdev
0 2 4
AnnualTemp
sdevlarge
0.0 2.5 5.0 7.5 10.0
Aridity
avg
0 3000 6000 9000 12000
Aridity
sdev
0 1000 2000 3000
Aridity
sdevlarge
0 2000 4000 6000 8000
AvgMaxNDVI
avg
0.00 0.25 0.50 0.75
AvgMaxNDVI
sdev
0.0 0.1 0.2 0.3
AvgMaxNDVI
sdevlarge
0.0 0.1 0.2
Elevation
avg
0 1000 2000 3000 4000 5000
Elevation
sdev
0 250 500 750 1000
Elevation
sdevlarge
0 500 1000 1500
MeanDiurnalRange
avg
10 15 20
MeanDiurnalRange
sdev
0.0 0.5 1.0 1.5 2.0 2.5
MeanDiurnalRange
sdevlarge
0 1 2 3
Pet
avg
1000 2000 3000 4000
Pet
sdev
0 100 200 300 400
Pet
sdevlarge
0 100 200 300 400 500
PrecDriestMonth
avg
0 20 40
PrecDriestMonth
sdev
0 3 6 9
PrecDriestMonth
sdevlarge
0 5 10 15 20
0 25 50 75
richness
value

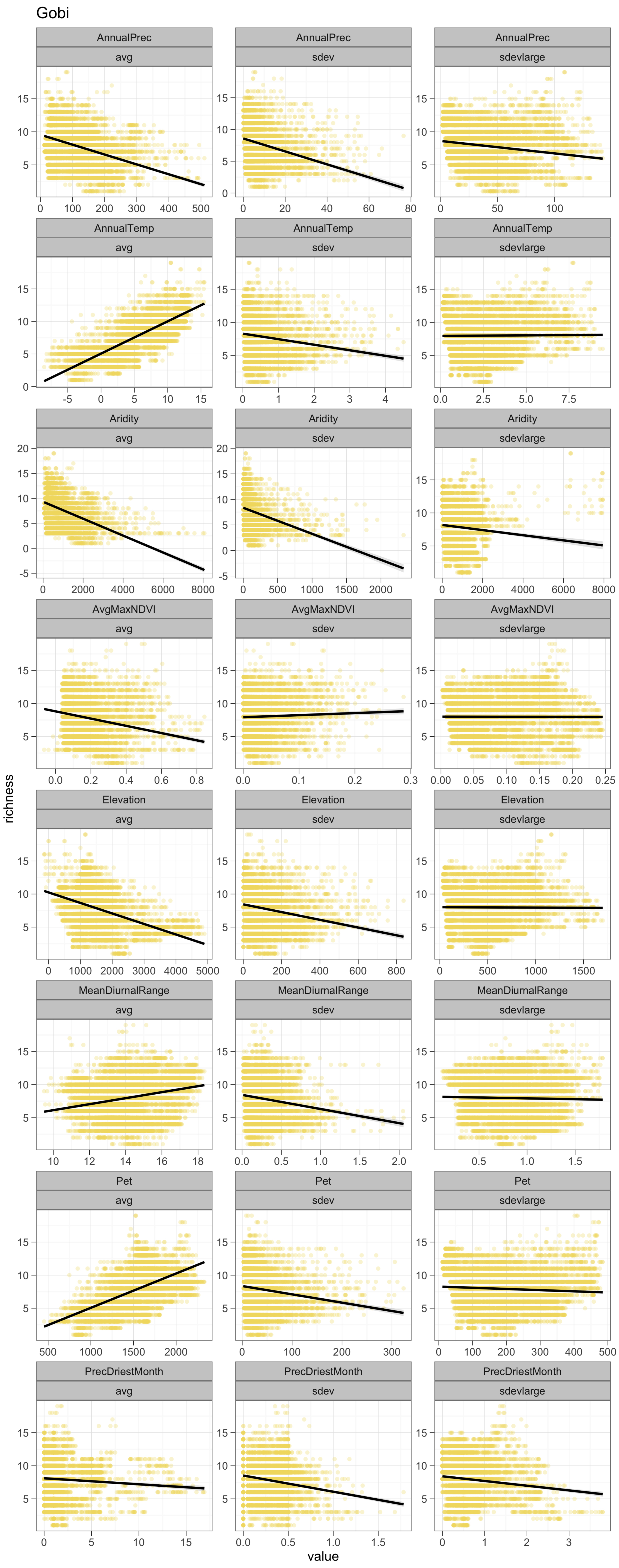
Gobi
AnnualPrec
avg
AnnualPrec
sdev
AnnualPrec
sdevlarge
AnnualTemp
avg
AnnualTemp
sdev
AnnualTemp
sdevlarge
Aridity
avg
Aridity
sdev
Aridity
sdevlarge
AvgMaxNDVI
avg
AvgMaxNDVI
sdev
AvgMaxNDVI
sdevlarge
Elevation
avg
Elevation
sdev
Elevation
sdevlarge
MeanDiurnalRange
avg
MeanDiurnalRange
sdev
MeanDiurnalRange
sdevlarge
Pet
avg
Pet
sdev
Pet
sdevlarge
PrecDriestMonth
avg
PrecDriestMonth
sdev
PrecDriestMonth
sdevlarge
richness
value

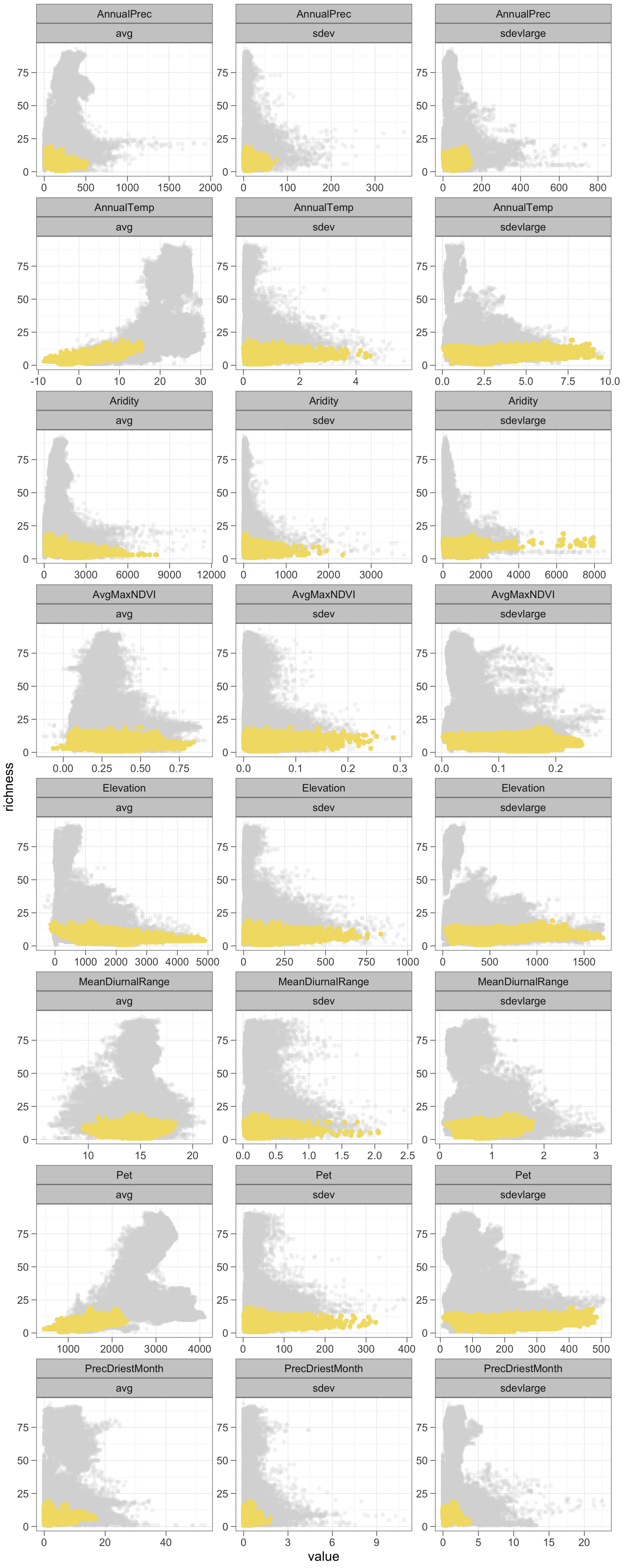
AnnualPrec
avg
sdev
sdevlarge
AnnualTemp
Aridity
AvgMaxNDVI
Elevation
MeanDiurnalRange
Pet
PrecDriestMonth
richness
value

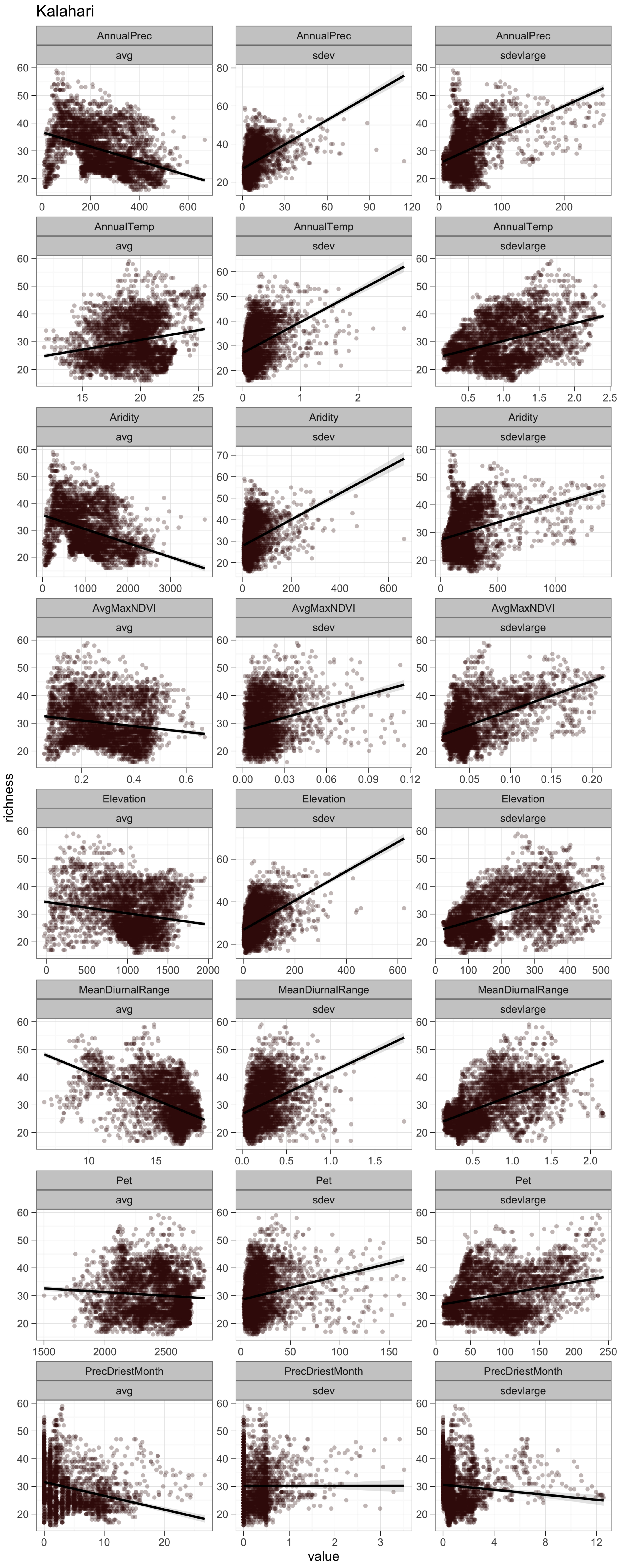

Kalahari
AnnualPrec avg
AnnualPrec sdev
AnnualPrec sdevlarge
AnnualTemp avg
AnnualTemp sdev
AnnualTemp sdevlarge
Aridity avg
Aridity sdev
Aridity sdevlarge
AvgMaxNDVI avg
AvgMaxNDVI sdev
AvgMaxNDVI sdevlarge
Elevation avg
Elevation sdev
Elevation sdevlarge
MeanDiurnalRange avg
MeanDiurnalRange sdev
MeanDiurnalRange sdevlarge
Pet avg
Pet sdev
Pet sdevlarge
PrecDriestMonth avg
PrecDriestMonth sdev
PrecDriestMonth sdevlarge
richness
value

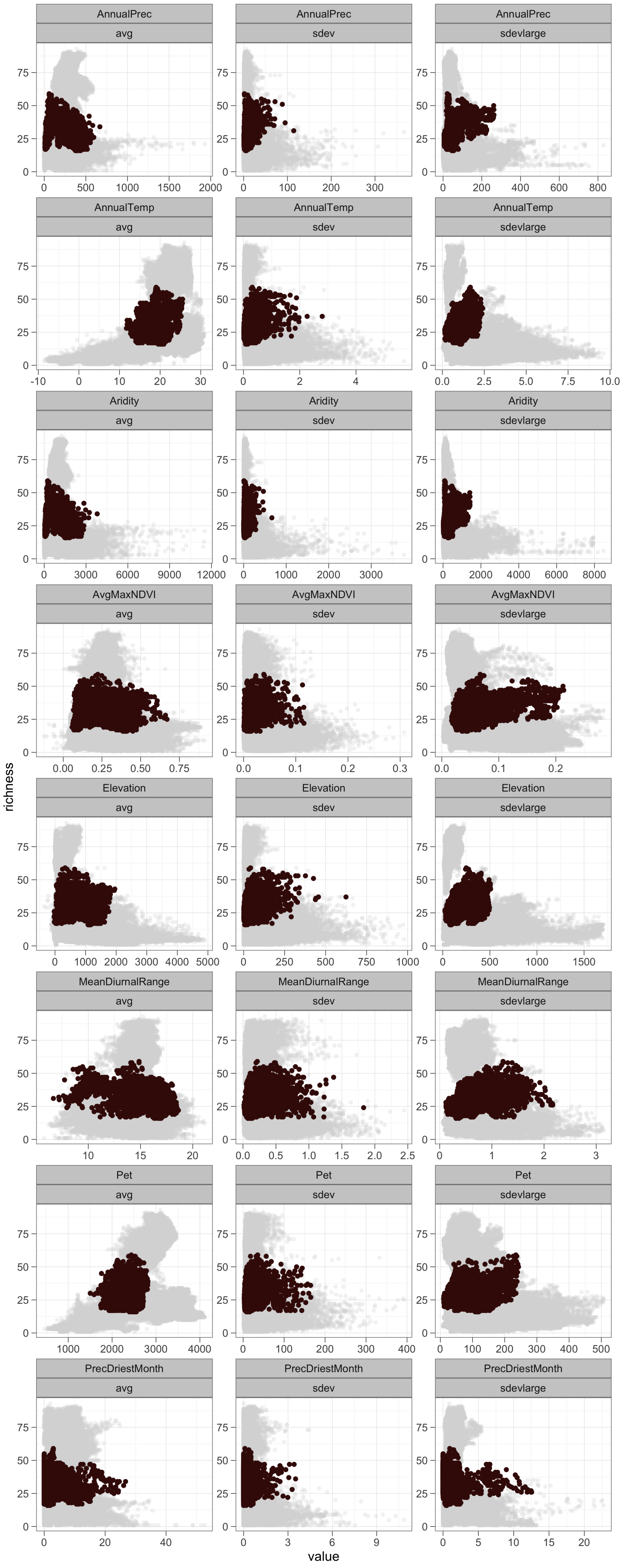
AnnualPrec
avg
sdev
sdevlarge
AnnualTemp
Aridity
AvgMaxNDVI
Elevation
MeanDiurnalRange
Pet
PrecDriestMonth
richness
value

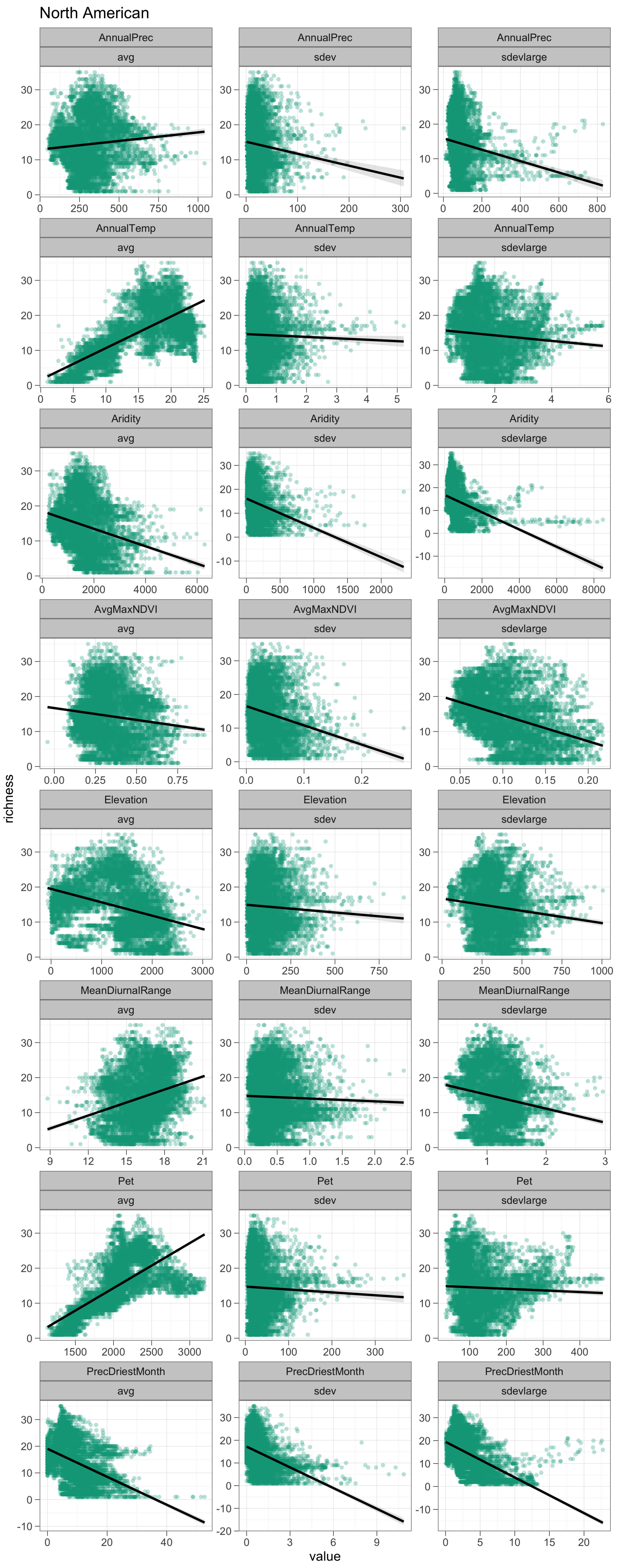
North American
AnnualPrec
avg
AnnualPrec
sdev
AnnualPrec
sdevlarge
AnnualTemp
avg
AnnualTemp
sdev
AnnualTemp
sdevlarge
Aridity
avg
Aridity
sdev
Aridity
sdevlarge
AvgMaxNDVI
avg
AvgMaxNDVI
sdev
AvgMaxNDVI
sdevlarge
Elevation
avg
Elevation
sdev
Elevation
sdevlarge
MeanDiurnalRange
avg
MeanDiurnalRange
sdev
MeanDiurnalRange
sdevlarge
Pet
avg
Pet
sdev
Pet
sdevlarge
PrecDriestMonth
avg
PrecDriestMonth
sdev
PrecDriestMonth
sdevlarge
0 250 500 750 1000
0 100 200 300
0 200 400 600 800
0 5 10 15 20 25
0 1 2 3 4 5
2 4 6
2000 4000 6000
0 500 1000 1500 2000
0 2000 4000 6000 8000
0.00 0.25 0.50 0.75
0.0 0.1 0.2
0.05 0.10 0.15 0.20
0 1000 2000 3000
0 250 500 750
0 250 500 750 1000
9 12 15 18 21
0.0 0.5 1.0 1.5 2.0 2.5
1 2 3
1500 2000 2500 3000
0 100 200 300
100 200 300 400
0 20 40
0 3 6 9
0 5 10 15 20
richness
value

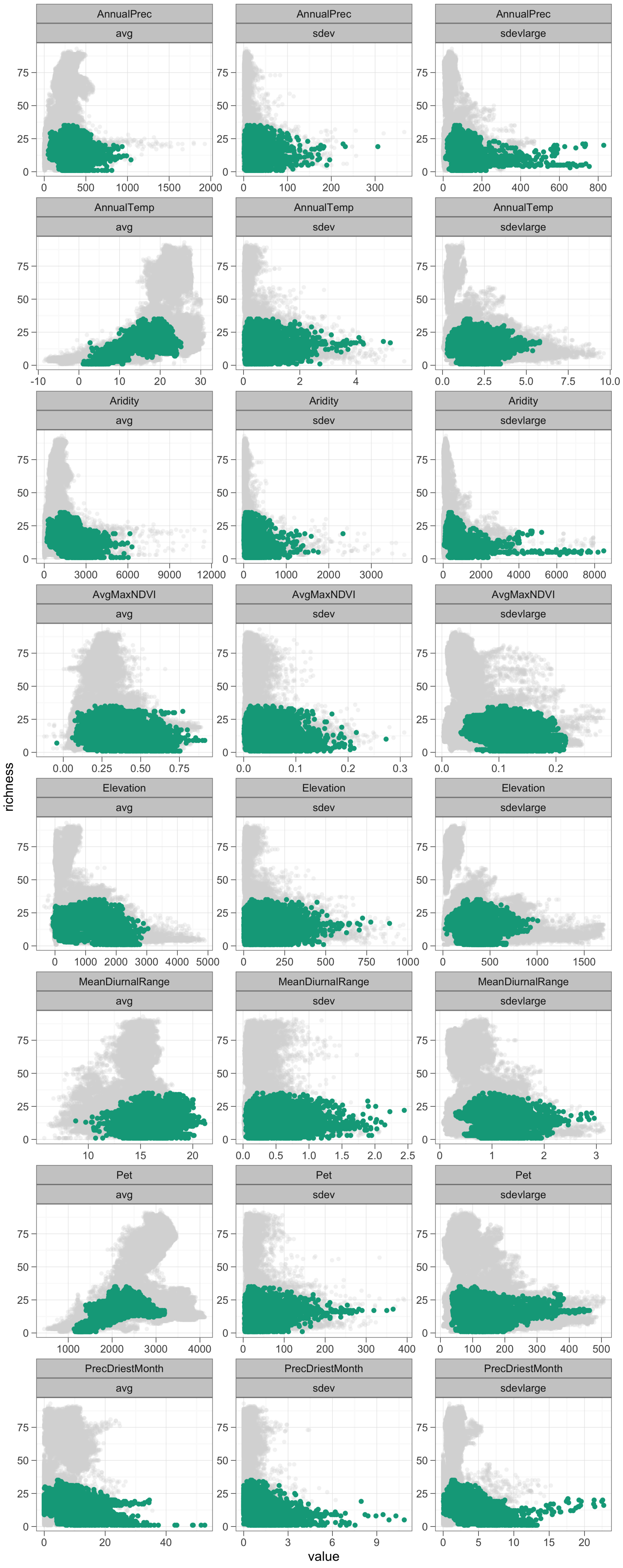

AnnualPrec
avg
sdev
sdevlarge
AnnualTemp
Aridity
AvgMaxNDVI
Elevation
MeanDiurnalRange
Pet
PrecDriestMonth
richness
value

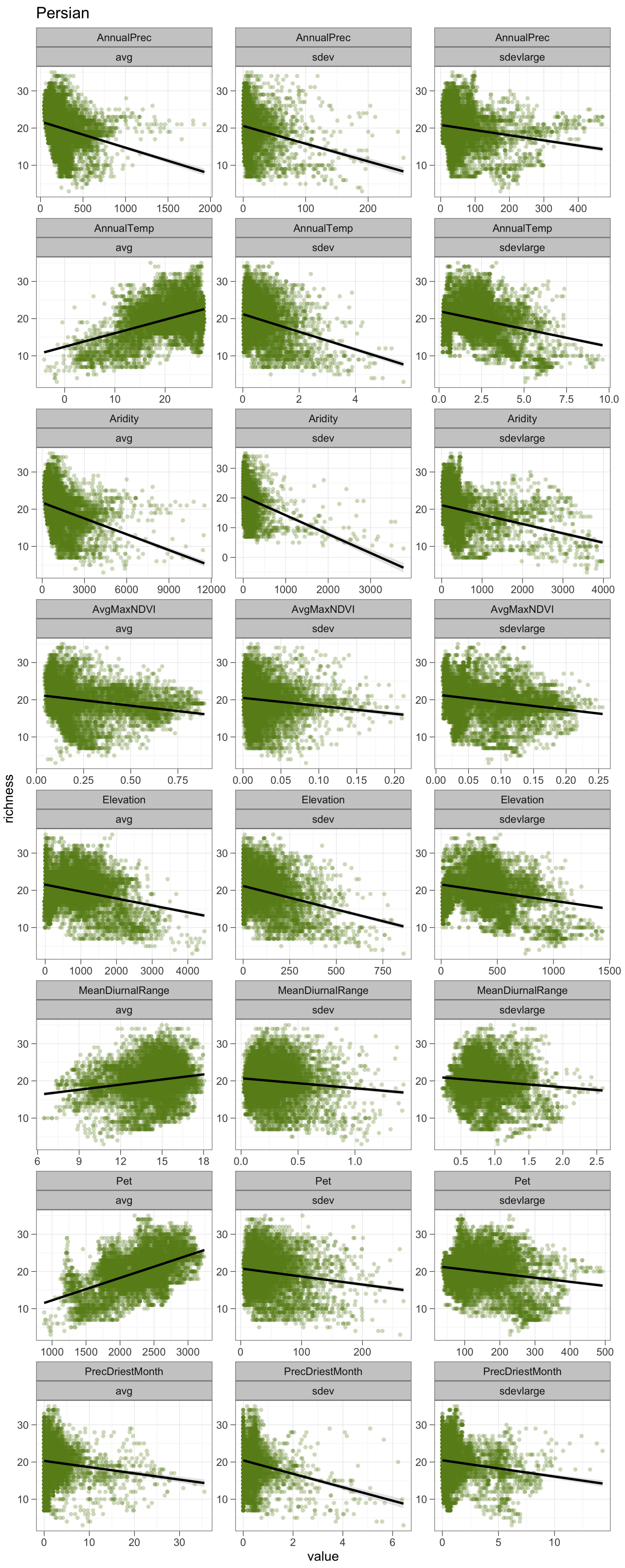

Persian
AnnualPrec
avg
sdev
sdevlarge
AnnualTemp
Aridity
AvgMaxNDVI
Elevation
MeanDiurnalRange
Pet
PrecDriestMonth
richness
value

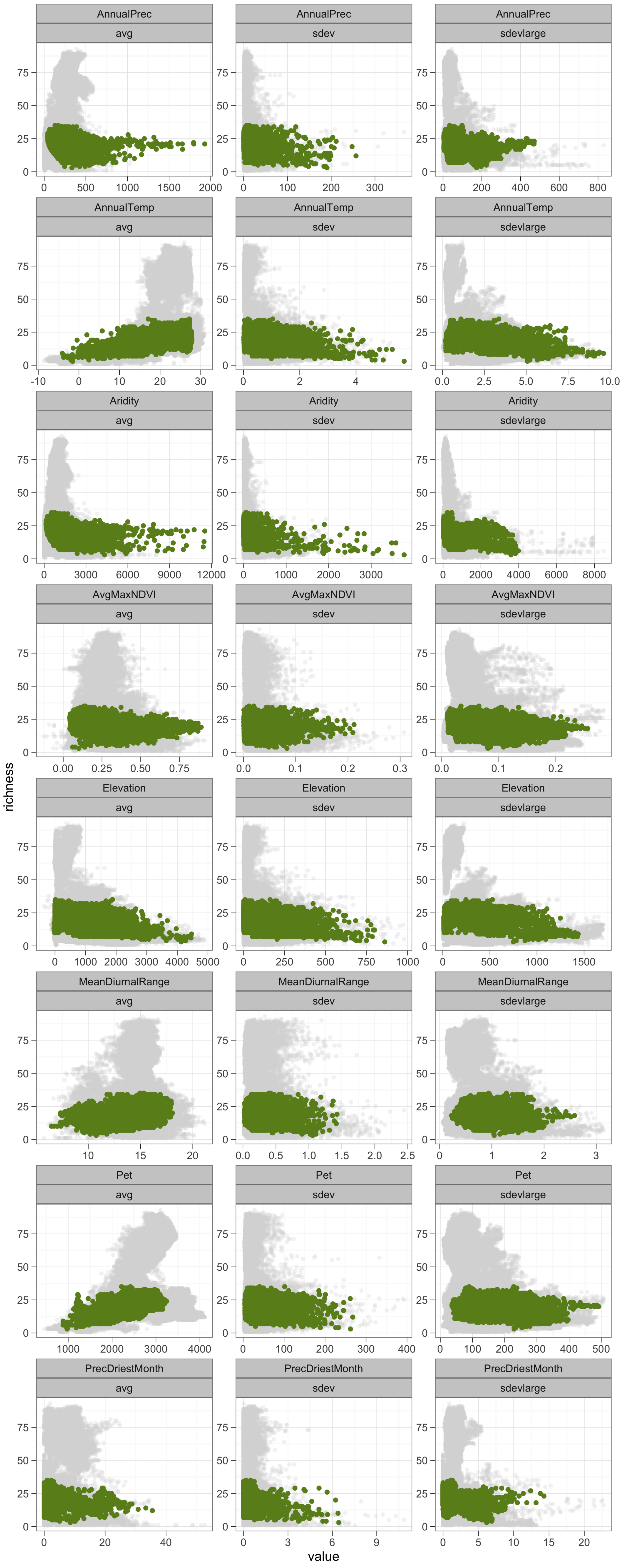

AnnualPrec
avg
sdev
sdevlarge
AnnualTemp
Aridity
AvgMaxNDVI
Elevation
MeanDiurnalRange
Pet
PrecDriestMonth
richness
value

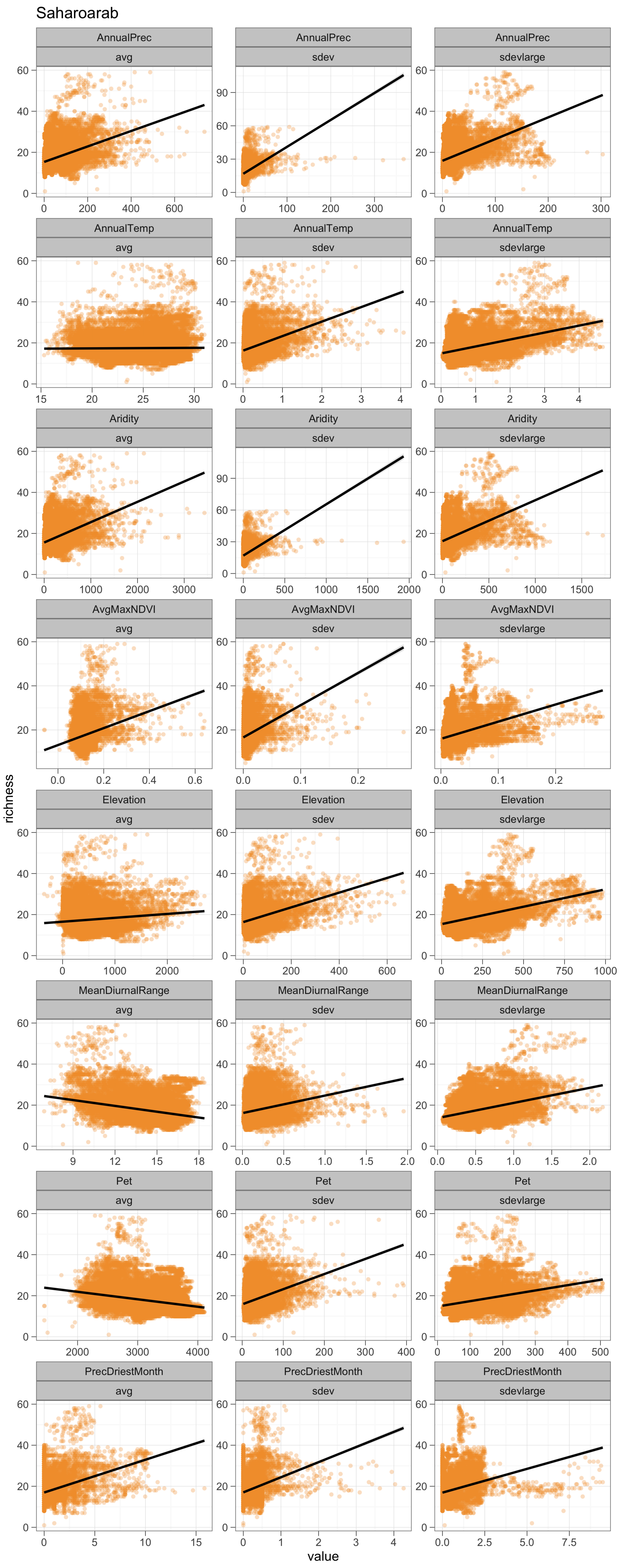

Saharoarab
AnnualPrec avg
AnnualPrec sdev
AnnualPrec sdevlarge
AnnualTemp avg
AnnualTemp sdev
AnnualTemp sdevlarge
Aridity avg
Aridity sdev
Aridity sdevlarge
AvgMaxNDVI avg
AvgMaxNDVI sdev
AvgMaxNDVI sdevlarge
Elevation avg
Elevation sdev
Elevation sdevlarge
MeanDiurnalRange avg
MeanDiurnalRange sdev
MeanDiurnalRange sdevlarge
Pet avg
Pet sdev
Pet sdevlarge
PrecDriestMonth avg
PrecDriestMonth sdev
PrecDriestMonth sdevlarge
richness
value

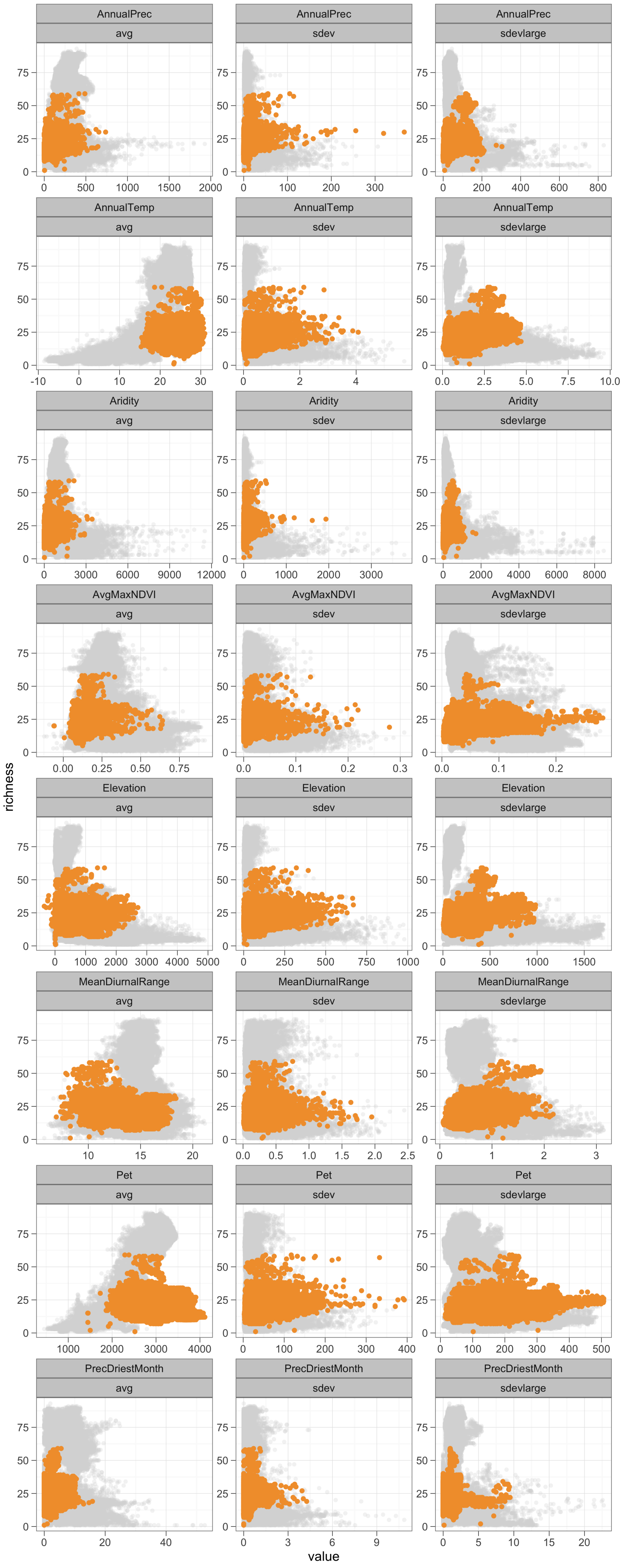
AnnualPrec
avg
sdev
sdevlarge
AnnualTemp
Aridity
AvgMaxNDVI
Elevation
MeanDiurnalRange
Pet
PrecDriestMonth
richness
value
