## Supplementary figures and images for "Desert lizard diversity worldwide: effects of environment, time, and evolutionary rate"

### Fig. S3

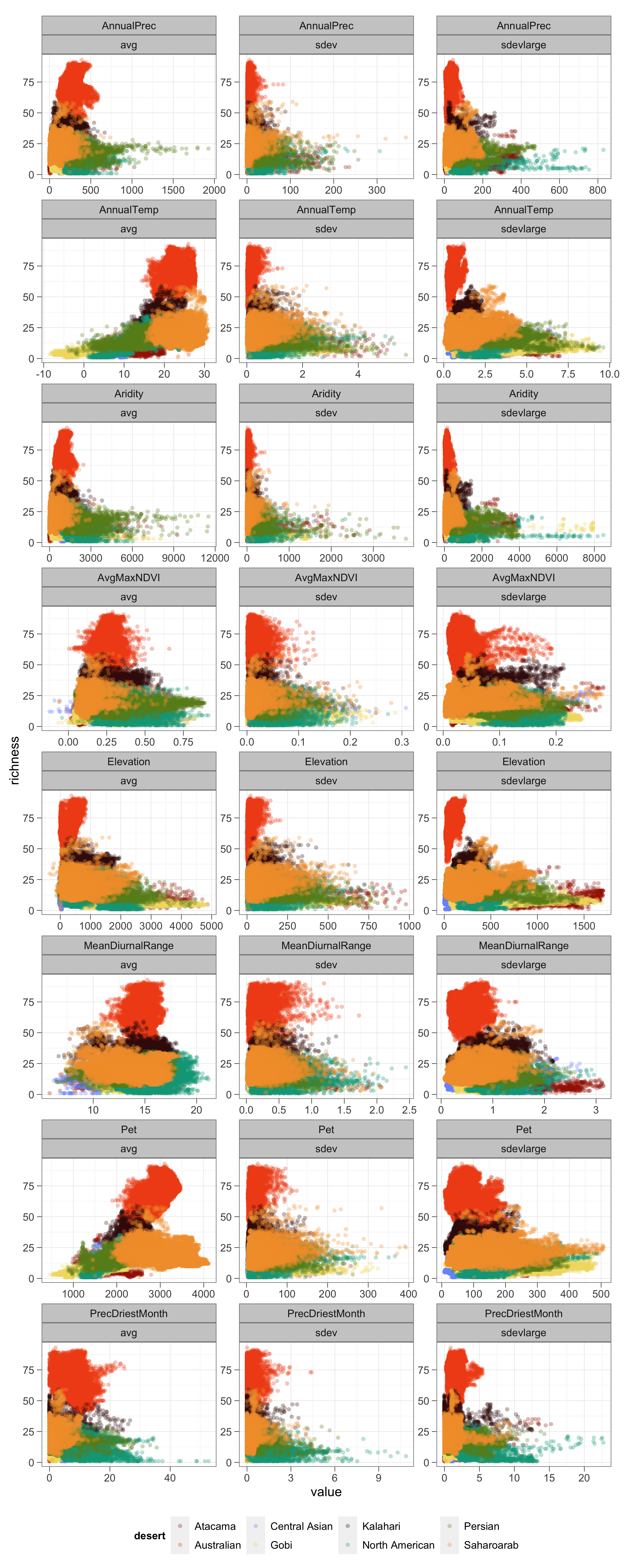
